# Accuracy and Reproducibility of Somatic Point Mutation Calling in Clinical-Type Targeted Sequencing Data

**DOI:** 10.1101/2019.12.31.891952

**Authors:** Ali Karimnezhad, Gareth A. Palidwor, Kednapa Thavorn, David J. Stewart, Pearl A. Campbell, Bryan Lo, Theodore J. Perkins

## Abstract

**Background:** Treating cancer depends in part on identifying the mutations driving each patient’s disease. Many clinical laboratories are adopting high-throughput sequencing for assaying patients’ tumours, applying targeted panels to formalin-fixed paraffin-embedded tumour tissues to detect clinically-relevant mutations. While there have been some benchmarking and best practices studies of this scenario, much variant-calling work focuses on whole-genome or whole-exome studies, with fresh or fresh-frozen tissue. Thus, definitive guidance on best choices for sequencing platforms, sequencing strategies, and variant calling for clinical variant detection is still being developed.

**Results:** Because ground truth for clinical specimens is rarely known, we used the well-characterized Coriell cell lines GM12878 and GM12877 to generate data. We prepared samples to mimic as closely as possible clinical biopsies, including formalin fixation and paraffin embedding. We evaluated two well-known targeted sequencing panels, Illumina’s TruSight 170 panel and the Oncomine Focus panel. Sequencing was performed on an Illumina NextSeq500 and an Ion Torrent PGM respectively. We performed multiple biological replicates of each assay, to test reproducibility. Finally, we applied five different public and freely-available somatic single-nucleotide variant (SNV) callers to the data, MuTect2, SAMtools, VarScan2, Pisces and VarDict. Although the TruSight 170 and Oncomine Focus panels cover different amounts of the genome, we did not observe major differences in variant calling success within the regions that each covers. We observed substantial discrepancies between the five variant callers. All had high sensitivity, detecting known SNVs, but highly varying and non-overlapping false positive detections. Harmonizing variant caller parameters or intersecting the results of multiple variant callers reduced disagreements. However, intersecting results from biological replicates was even better at eliminating false positives.

**Conclusions:** Reproducibility and accuracy of targeted clinical sequencing results depends less on sequencing platform and panel than on downstream bioinformatics and biological variability. Differences in variant callers’ default parameters are a greater influence on algorithm disagreement than other differences between the algorithms. Contrary to typical clinical practice, we recommend analyzing replicate samples, as this greatly decreases false positive calls.

## Background

Next generation sequencing (NGS) technologies have been used to catalogue genetic mutations in cancer [1]. Studies employing NGS have identified specific genetic mutations that reliably predict therapeutic success with targeted treatment regimens in many forms of cancer [2], including non-small cell lung cancer (NSCLC), which is our long-term focus. Importantly, patients with oncogenic driver mutations have better tumour control with targeted agents than with chemotherapy, while those lacking such a mutation derive more benefit from chemotherapy [3]. Thus, accurate identification of the mutated or non-mutated status of key genomic sites is critical for patient therapy.

Developments in NGS technologies are empowering the analyses of whole cancer genomes, providing insights into the task of somatic mutation calling [4] and have made it possible to characterize the genomic alterations in a tumour in an unbiased manner [5]. With the advancement of NGS technologies, the number of large-scale projects (especially cancer projects) dealing with somatic point mutation in various tumour types has been increasing rapidly. However, in clinical practice, there is only a limited number of actionable mutations that are of interest—those for which a specific therapy is recommended, or perhaps for which a clinical trial may be ongoing. In such cases, targeted sequencing is the preferred option, offering lower cost and higher coverage of areas of interest [6].

The majority of mutation callers available in the literature have been designed to analyze matched tumour-normal samples [7], as comparing the two helps discriminate cancer-related and non-cancer-related mutations. However, many clinical labs do not routinely acquire matched healthy tissue, so that variant calling must be performed on the tumour tissue only. As such, tumour-only mutation callers have also been developed. A few mutation callers such as Mutect2 and VarDict are versatile enough to analyze both matched tumour-normal and tumour-only samples. For a recent up-to-date list of mutation callers, readers may refer to [7], where 46 programs are reviewed. In this study, we focus on tumour-only mutation calling.

A typical workflow of mutation calling can be divided into three steps. First, reads are processed and low-quality bases and any exogenous sequences such as sequencing adapters are excluded from the reads. This can be performed by using tools such as Cutadapt [8] and NGS QC Toolkit [9]. Second, the cleaned reads are mapped to a reference genome. This base-to-base alignment can be done by using common tools such as BWA aligner [10] for DNA sequencing or TopHat [11] for RNA sequencing. The last step of the process is to separate real mutations from artifacts that might be present due to a process of library preparation, read errors, mapping errors, and so on. Some mutation callers perform the above three-step process as a built-in procedure (see [7]), while others start from the point of cleaned, mapped reads.

Several analysis packages with different algorithms have been introduced in recent years to increase mutation detection accuracy. Significant discrepancies between the results of different algorithms have been observed, which leads to a difficulty in selecting candidate mutations for validation [12]. Such disagreement seems to partially root from different error models and assumptions made in each algorithm. In addition, various sources of errors such as sequencing errors and read alignment errors make the process more challenging. Thus, detection accuracy remains questionable and in fact, has become a major challenge.

Previous benchmarking studies offer some guidance in this regard, each having its strengths and weaknesses. For example, Spencer et al. [13] evaluated four mutation callers on a compendium of data from several cell lines and mixtures of DNA from those cell lines—artificially “creating” mutations at different allele frequencies. However, they focused on only one targeting panel, the Washington University comprehensive cancer set WUCaMP27 version 1.0, and one sequencing platform, the Illumina HiSeq 2000. Also problematically, they defined gold standard mutations by intersecting variant calls from GATK and SAMtools, two of the four programs they benchmarked. Derryberry et al. [14] focused on the technical reproducibility of variant calls, studying 55 replicate pairs of data from gliobastoma tumours. However, this was whole genome sequencing data, where coverage was substantially lower than for targeted panels. Xu et al. [15] performed a thorough comparison of mutation callers on exome sequencing data of pure and admixed cell lines, where gold-standard mutations where established by prior independent work. Moreover, they explored, to some extent, the effects of varying variant-calling thresholds. However, their work was limited to tumour-normal variant calling. Numerous other benchmarking studies have been performed; we mention these only as examples. Of note, new “best practices” for benchmarking variant calling have recently been proposed [16].

Of course, no single study can exhaustively address all possible relevant issues in variant calling. We seek here to add to the conversation by comparing five variant callers, on data from two cell lines, sequenced on two different platforms using two different targeted panels, in multiple biological replicates. We minimize circularity in defining gold-standard mutations by relying on previous high-quality work using independent data and variant calling methods. We study agreements and disagreements in variant calls, depending on algorithms, algorithm settings, and between replicates. Our key findings are: that sequencing platforms are not a major influence on variant calling; that variant callers can disagree wildly when used with default/recommended settings, but that they agree much more when settings are harmonized; and that intersecting the results from different algorithms (which is not unusual in research practice, though possibly unusual in clinical practice) or from replicate samples (which is definitely unusual in clinical practice), can greatly reduce false positive calls while maintaining sensitivity for detecting genuine mutations.

## Results and discussion

For the purpose of benchmarking different variant callers, we used clinical sample handling protocols to perform targeted DNA or DNA/RNA sequencing on multiple biological replicates of multiple cell lines. Because accuracy assessment requires us to know which mutations should be present in the samples, we selected two very well characterized Coriel Cell lines, GM12878 and GM12877. As our long-term goal is understanding best methods for variant calling in formalin-fixed, paraffin-embedded (FFPE) lung cancer fine-needle aspirates (FNAs), we performed independent preparations of the GM12878 and GM128977 cells for analysis in FFPE cytoblocks. We selected two targeted assays for testing. One was the TruSight170 Tumor Assay (TST170) from Illumina. TST170 is a comprehensive RNA/DNA hybrid-capture assay that provides full coverage for 170 solid tumor-associated genes. It is amenable to testing on FFPE samples of relatively limited sample availability. This assay tests for the presence of multiple classes of structural variants that are relevant to clinical diagnostics including SNVs, fusions, copy number variation, and INDELs. Here, however, we will focus only on the SNVs. The TST170 sequencing was performed on an Illumina NextSeq500 sequencer. We also tested the Oncomine Focus panel, which again is amenable to FFPE samples with limited material. Oncomine Focus covers 47 genes, and sequencing was performed on an Ion Torrent PGM. The sequencing data is available through SRA under accession PRJNA614006. Table 1 summarizes several key features of the sequencing data we generated. For the TST170 panel, read depths varied from approximately 2.4 million to over 13 million per sample. Within the genomic regions covered by the panel, this resulted in average coverage levels ranging from 203 to over 1000 reads per basepair.

**Table 1:** Summary of read depth and coverage (average reads covering each basepair covered by the panel) for TruSight Tumor 170 (TST170) and Oncomine Focus (OF) panels.

|  | TST170 - 12878 |  | TST170 - 12877 |  | OF - 12878 |  | OF - 12877 |  |
| --- | --- | --- | --- | --- | --- | --- | --- | --- |
| Rep | Reads | Cover | Reads | Cover | Reads | Cover | Reads | Cover |
| 1 | 13275831 | 1056 | 3526629 | 300 | 835719 | 2437 | 602628 | 1829 |
| 2 | 7598224 | 659 | 3285901 | 280 | 213469 | 625 | 769173 | 2252 |
| 3 | 8279644 | 725 | 2440647 | 203 | 138350 | 415 | 1566916 | 4338 |
| 4 | 7435784 | 640 | 3640546 | 323 | — | — | — | — |
| 5 | 7519631 | 648 | 4731704 | 412 | — | — | — | — |
| 6 | 7775631 | 672 | 4101646 | 366 | — | — | — | — |

For the Oncomine data, read depths ranged from 1.3 million to over 15 million, resulting in coverages ranging from 415 to over 4000 reads per basepair.

### Sequencing data shows expected SNVs in GM12878 and GM12877 cell lines

Recent work has established a comprehensive and genome-wide catalog of high-confidence variants for a collection of Coriell cell lines, including GM12878 and GM12877 [17]. That work relied on whole-genome sequencing data of 17 individuals in a three-generation pedigree, and variant calling using four programs: FreeBayes [18], Platypus [19], GATK3 [20] and Strelka [21]. By intersecting the locations of these known mutations with the TST170 genomic regions, we determined that our GM12878 and GM12877 data should contain 343 and 336 of these mutations, respectively. Intersecting with the Oncomine Focus genomic regions, we predicted that 26 and 24 known mutations should appear in GM12878 and GM12877 data respectively.

The first thing we checked was whether there was indeed evidence for these high-confidence, gold-standard mutations in our data. While the ability to call a variant depends on many factors, two key factors are the coverage (or read depth) at the site, and the frequency of the alternative (i.e. non-reference) allele (allele frequency = AF). We mapped all read data to the hg19 genome (see Methods for details), and computed the read depths and empirical allele frequencies for the known mutations for both cell lines and both panels using Bam-readcount (https://github.com/genome/bam-readcount). Figure 1 shows the results.

**Figure 1:**
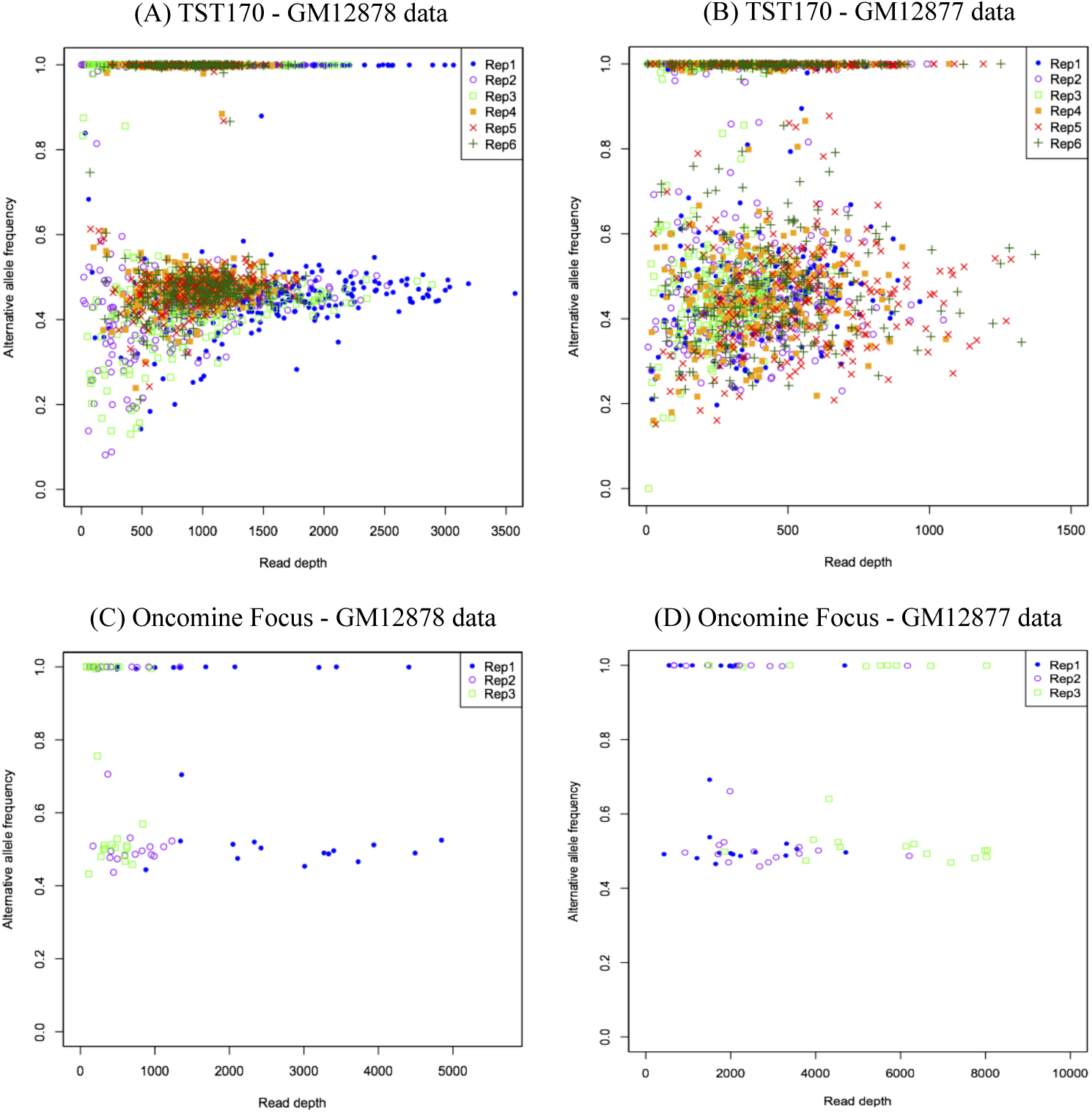
Distribution of alternative allele frequency and read depth of known gold-standard mutations in the data.

Figure 1A, for example, shows AF and read depth for the gold-standard mutations in all six replicates of our TST170 data from the GM12878 cell line, with the mutations in each replicate denoted by a different color and shape of mark. Several points are worth nothing. First, regardless of replicate, there are two main clusters of points. One cluster is at or very near AF = 1, representing homozygous mutations relative to hg19. The other cluster centers near AF = 0.5, but with substantially greater spread, and represents heterozygous mutations. As would be expected on simple random-sampling grounds, there is more spread in empirical AF around AF = 0.5 when read depth is lower, with some heterozgyous variants having AF near 0.1 and others over 0.8. In all six replicates, there is no known mutation with zero read depth nor zero alternative AF. Thus we can say that every dataset contains at least some evidence for every known gold-standard mutation.

Figure 1B shows the same type of plot for our TST170 GM12877 data. The patterns of AF and depth are similar, although we observe several qualitative differences from Figure 1A. First, the spread in empirical AF for the heterozygous mutations is greater. We suspect this is due in part to the lower overall read depths of these experiments, which came out at roughly half compared to GM12878. Few mutations exceed a coverage of 1000 reads in our GM12877 data, whereas that is the approximate median for our GM12878 data. It is not clear if this is the only factor leading to the greater AF spread. We also observe slightly greater frequency of mutations just below the level of AF=1. This may suggest a slightly higher per-base read error rate or possibly alignment errors. Excepting one mutation in replicate 3, there is no known mutation with zero alternative AF.

Figures 1C-D show similar plots for the Oncomine Focus data sets. The plots are much sparser, because these panels cover so much less genomic territory and so many fewer mutations. But the same broad pattern emerges. We see the gold-standard mutations having observed alternative AFs either near 0.5 or 1. In our Oncomine Focus data, there is greater difference in total read depth between replicates, and so unsurprisingly there is greater disparity in read depths at the mutation sites between replicates. Again, we see that every mutation is supported by at least some evidence. Collectively then, with the exception of just a single site in one TST170 replicate of GM12877 data, our data contains evidence of all the expected gold-standard mutations.

The converse question is whether our data contain evidence of other mutations. These may be genuine mutations that have been missed by other variant calling efforts, or that arose spotaneously in our copies of the cell lines. Or they may be artifacts of some sort masquerading as genuine mutations. To study this question, we produced the same type of AF-versus-read-depth plots as in Figure 1, but for all the genomic positions covered by the panels that are not on the gold-standard list of mutations. The results can be seen in Figure 2. Panel A in particular shows the results for all six replicates of GM12878 using the TST170 panel. Unlike in Figure 1, where we saw substantial clouds of points suggesting heterozygous and homozygous mutations, the vast majority of these positions hover around low allele frequency. If these are non-mutated positions, then ideally the AF should read as zero; or, allowing for a small per-base read error rate below 1%, small non-zero AFs could be expected. We see a substantial number of positions with AF at 0.2 or higher, even at relatively high coverage like 1000 reads or more. The fact that the apparent distribution of AFs gravitates towards zero as coverage increases suggests these higher AFs may be errors of some kind. Regardless, some of these positions have the potential to be confused with the genuine, gold-standard mutations. Comparing Figures 1A and 2A, we can see that in terms of AF and coverage, there is some overlap between gold-standard and presumed non-mutated sites. The data in Figure 2A also suggests a small number of sites that may be genuine mutations—sites where coverage is high (at least 500bp) and AF is also high (0.5 or more). From this plot, one cannot tell whether different replicates support potential genuine mutations at the same or different sites. We will return to this question later, when we analyze agreement in mutation calls across replicates. Figure 2B-D shows similar results for the other three conditions.

**Figure 2:**
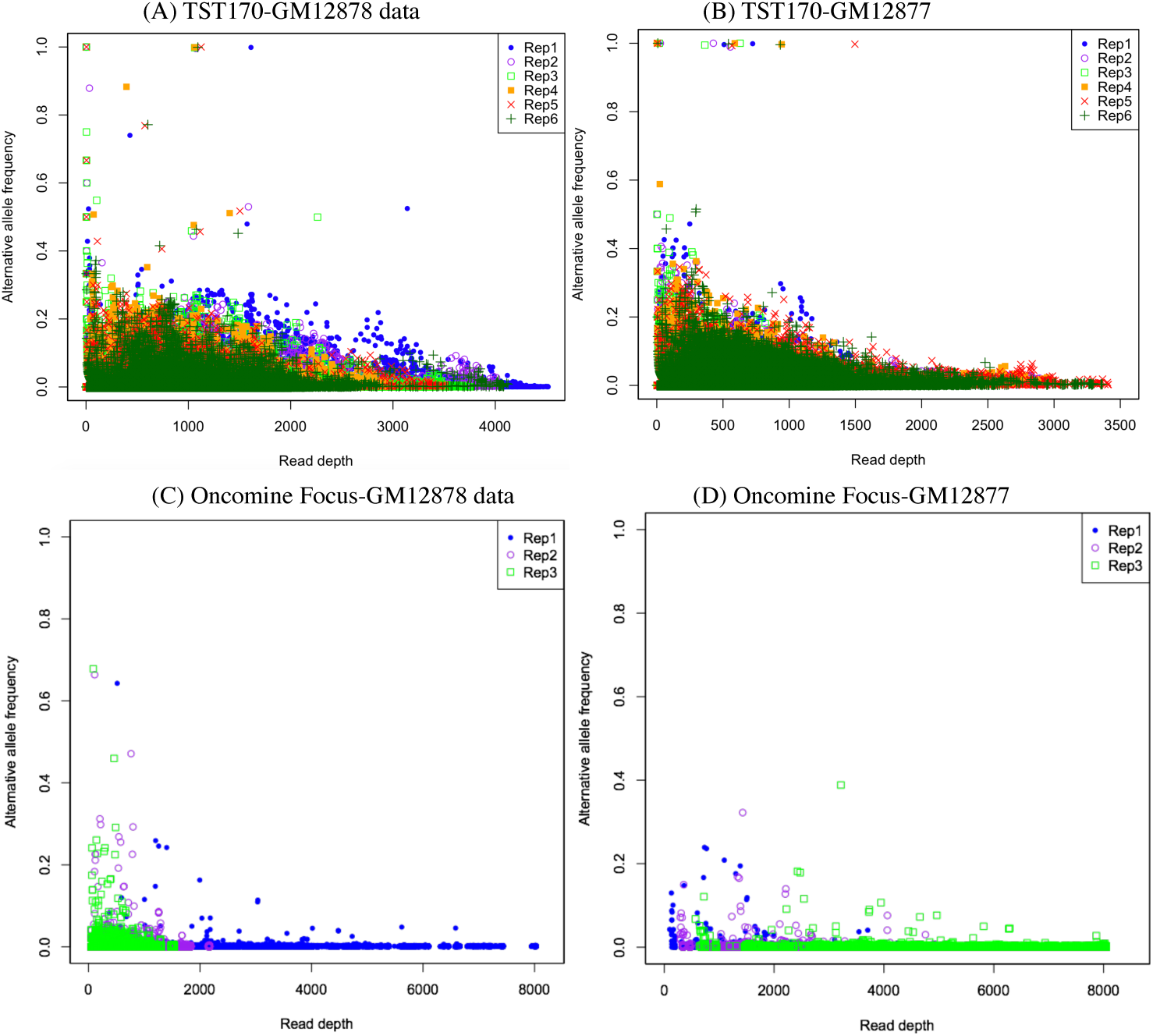
Distribution of alternative allele frequency and read depth at non-gold-standard mutation sites in the data.

### Variant callers disagree greatly under default configurations

We investigated the performance of five variant callers in separating the gold-standard mutations from the non-mutated sites: SAMtools [22], VarScan2 [23], MuTect2 [24], VarDict [25] and Pisces [26]. See Methods for exact version numbers and other details. These five callers were chosen for different reasons. The SAMtools package is one of the most highly cited programs in all of bioinformatics, and its variant-calling facilities in particular have been used widely. VarScan2 is another highly cited and well-established package for mutation calling. MuTect2 is the latest version of the MuTect program, which won a DREAM somatic genotyping contest [27]. VarDict is a more recent program and has facilities designed specifically for clinical-type sequencing protocols. Finally, we included Pisces as the manufacturer-recommended variant caller by Illumina, one of the two sequencing platforms that we employed. Importantly, all five programs also have several key properties that recommended them for our study: they are freely available to use; their code is open source, so we could install it on our local machines and compute cluster; they are are capable of variant calling in tumor-only mode; and all output VCF files containing SNV calls that can readily be compared to our gold-standard mutation list and to each other.

Some clinical sequencing centers have the expertise to carefully tune the bioinformatics tools they employ, but many simply use some established or recommended pipeline with default settings for all parameters and filters. We first compared the variant callers under default settings, as downloaded and/or recommended in their associated publications (see Methods for details). Figure 3 summarizes the results of applying the five variant callers to each of the TST170 samples, while Figure 4 summarizes the results of applying the five variant callers to each of the Oncomine Focus samples. The first and second row of the figures refer to the GM12878 and GM12877 results, respectively.

**Figure 3:**
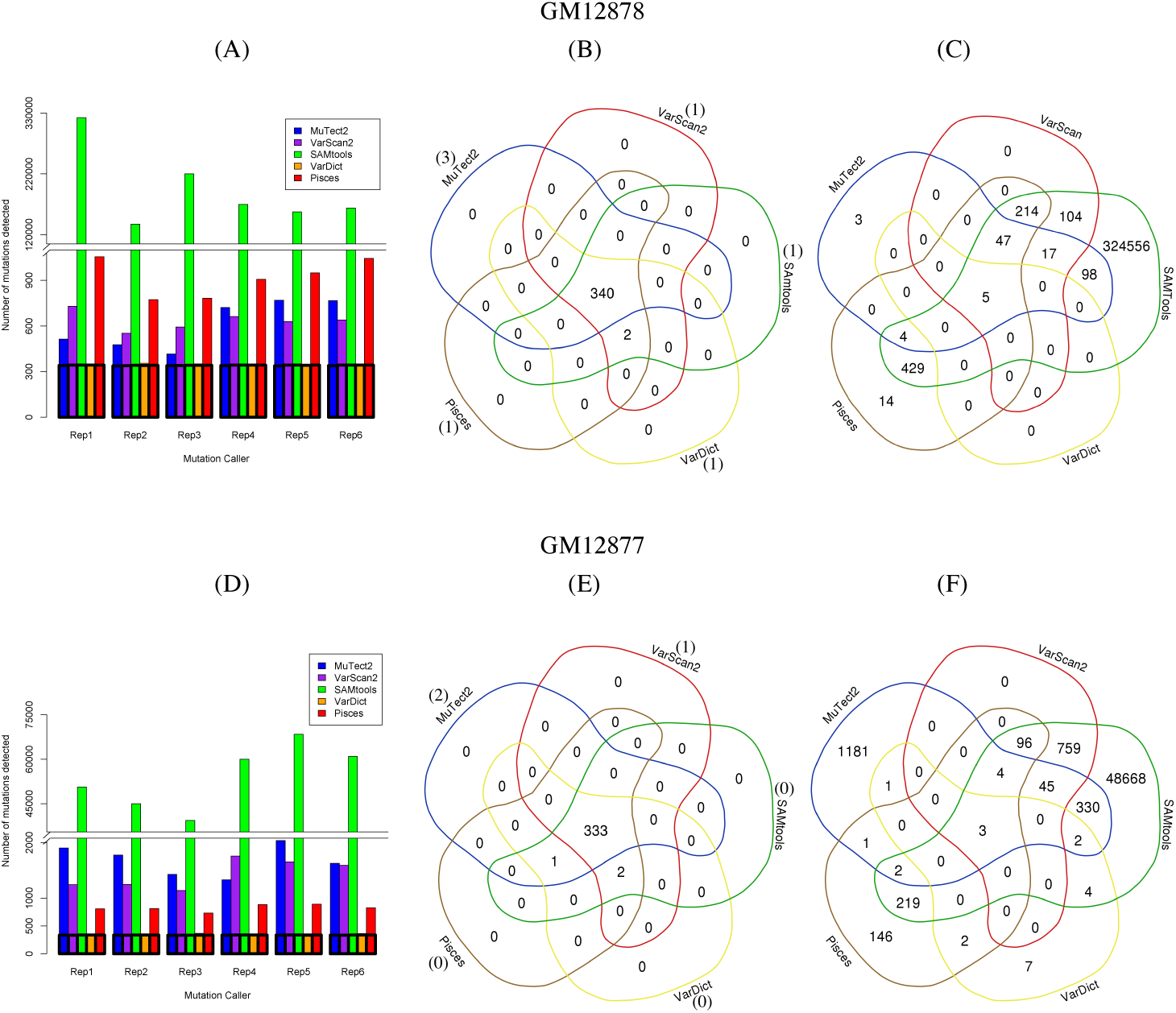
Comparison of five SNV calling approaches on TST 170 data sets using default parameters. (A,D) Numbers of variants called by each algorithm for GM12878 and GM12877 data respectively. (B,E) True positive and detections by the different algorithms on replicate 1 on GM12878 and GM12877. The numbers in parentheses represent total false negatives for each variant caller, i.e., the number of true variants not detected by the corresponding algorithm. (C,F) False positive calls by the different algorithms on replicate 1. Diagrams in this and subsequent figures made using the online tool http://bioinformatics.psb.ugent.be/webtools/Venn/.

**Figure 4:**
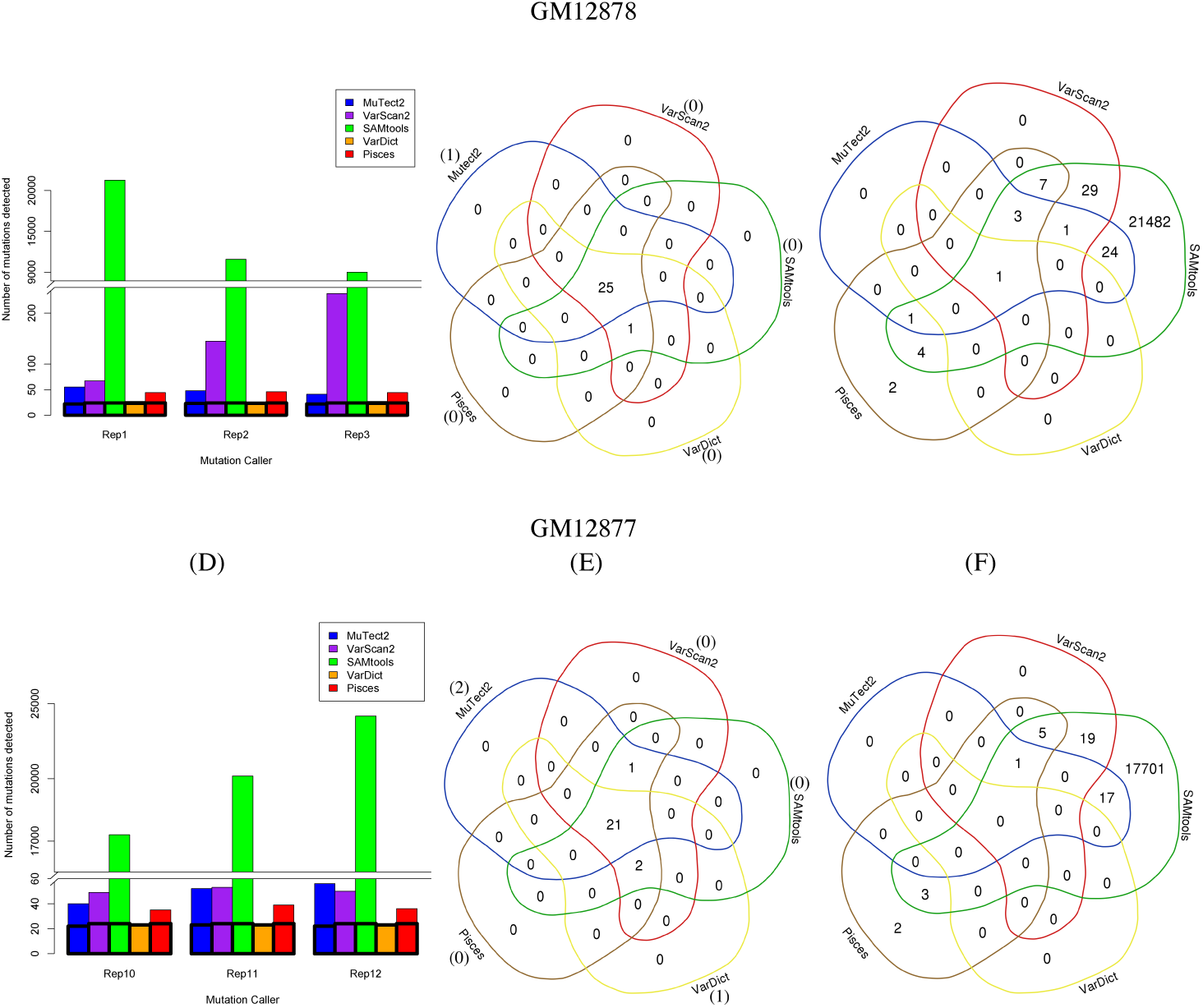
Comparison of five SNV calling approaches on Oncomine Focus data sets using default parameters. A,D) Numbers of variants called by each algorithm for GM12878 and GM12877 data respectively. (B,E) True positive and detections by the different algorithms on replicate 1 on GM12878 and GM12877. The numbers in parentheses represent total false negatives for each variant caller. (C,F) False positive calls by the different algorithms on replicate 1.

The barplots in Figure 3A show the number of mutations detected by each of the five callers in each of the six replicates of TST170 data. The highlighted black bars inside the barplots refer to the number of true positives. An immediate observation is that the various programs call wildly different numbers of SNVs from the same datasets. The most extreme is SAMtools, which calls between 128000 and 325000 mutations per replicate, compared to 340-1000 for the other callers. Keeping in mind that the TST170 panel covers just over half a million bases, this is an astonishingly high call rate. This happens because SAMtools’s default behavior is more or less to report anything that has any chance of being a mutation. Even a single read with a non-reference basepair is enough to cause SAMtools to flag a site. (We do not intend to criticize SAMtools here, but merely to warn about its default behaviour.) With the depth of coverage in our data and even a relatively low per-base error rate, many positions end up with some non-reference reads. Pisces is the next most profligate caller, reporting on average approximately three times as many sites as in our gold-standard list. At the opposite end of the spectrum, VarDict appears quite strict, calling only a few more sites than expected from the gold standard.

The Venn diagrams in Figures 3B and C break down in more detail the true and fale positive calls of the variant callers, and their agreements with each other, for replicate 1 specifically. The numbers in parenthesis in Figure 3B denote the number of false negatives. The good news is that all five variant callers correctly call almost all of the gold-standard mutations. All five variant callers agree on 340 of the 343 gold-standard mutations, and two more gold-standard mutations are called by all programs except Mutect2. All five programs miss one of the gold-standard mutations. In panel C, we see that although SAMtools calls a very large number of false positives, there remains a small number of false positives called uniquely by other programs. Pisces calls SNVs at 14 sites not called by any other program, and Mutect2 calls three unique SNVs. VarScan2 does not call any unique false positives, although it shares 104 calls with SAMtools that are not called by any other program. VarDict reports only five false positives, and these five are also reported by every other program. In terms of empirical false discovery rate (EFDR), performance ranges from an excellent 1% for VarDict, to a poor 99.9% for SAMtools.

For GM12877 data (Figure 3D-F) we see similar patterns. In panel D, we see that SAMtools call many possible variants, although not quite as many as for the GM12878 data. VarDict is again the most stringent caller, and others are in between. Figure 3E shows that all five callers agree on 333 of the 336 gold standard mutations, and the remaining three are picked up by four of five callers (Mutect2 misses 2, and VarScan2 misses the other). In Figure 3F we see that, perhaps as a consequence of SAMtools reporting fewer false positives, other programs report more unique false positives. Mutect2 reports over 1000, and Pisces over 100. Even VarDict calls seven false positives not called by any other algorithm. For these data, EFDR ranges from 5% (VarDict) to 99.3% (SAMtools).

Figure 4 presents the analogous results for the Oncomine Focus data. Absolute numbers are generally smaller, because the panel covers fewer total basepairs and fewer gold-standard mutations than the TST170 panel. But qualitative results are very similar. SAMtools calls large numbers of variants, almost all of which are false positives. VarDict calls very few false positives. All algorithms correctly detect most gold-standard mutations: 25 of 26 for GM12878 replicate 1 (Figure 4B), and 21 of 24 for GM12877 replicate 1(Figure 4E). Of the four gold-standard mutations not fully agreed-upon across the two cell lines, MuTect2 is responsible for missing three and VarDict for missing one. VarDict had only one false positive over all, while the other algorithms had roughly as many false positives and true positives, if not more.

### Harmonizing parameters makes variant calls more similar

Although each variant caller relies on a distinct statistical model, they also filter both inputs and results in various ways. Differences in the default parameters of these filters are an obvious possible explanation for divergent variant calls. Thus, we re-analyzed both the TST170 and Oncomine Focus data with the same five algorithms but setting their filtering parameters to be as similar as possible (see Methods). We set the minimum alternative allele frequency for a called variant to 0.01, the minimum coverage to 10 reads, the minimum base call quality to 20, and the minimum mapping quality to 20. These are not especially restrictive parameter choices, aimed at eliminating false positives calls. Later, we will return to the question of whether more restrictive filtering can improve performance and/or agreement between callers. Here, the focus is on making the filtering criteria of the algorithms as similar as possible.

Figure 5 summarizes the results of SNV calling with the harmonized parameters for the TST170 data sets, and can be compared to Figure 3, where results for default parameters are reported. In panels 5A and 5D, we see that harmonizing parameters greatly reduces the number of variants called by SAMtools. On the GM12878 data, the numbers are roughly on par with the other callers. On the GM12877 data, the numbers are still higher than for other callers, but they are 10 times or more lower than before parameter harmonization. Those panels also show a substantial increase in the number of mutations being called by VarDict. Much of this is due to our lowering of VarDict’s alternative allele frequency threshold from the default value of 0.05 to our harmonized value of 0.01. In panels 5B and 5E we see that the five variant callers continue to correctly identify and agree on most of the gold-standard variants. For GM12878, all five callers identify 340 of the 343 variants, and all programs except MuTect2 identify two more. For GM12878 the story is similar, with 333 of 336 fully agreed upon, and two more identified by all programs except MuTect2. The landscape of false positive calls is much different after parameter harmonization, however, as seen in panels 5C and 5F. Of course, there are many fewer false positives, because there are fewer variants being called over all. There are also many more variants being called uniquely by one caller and none of the four others. We also see all five callers agreeing on a total of 10 false positives, across both the GM12878 and GM12877 replicate 1 data, whereas there was only one before parameter harmonization. For GM12878, Pisces has the highest empirical false discovery rate (EFDR) at 64%, while VarDict has the lowest, at 9%. For GM12877 data, SAMtools has the highest EFDR at 91%, while Pisces has the lowest, at 50%.

**Figure 5:**
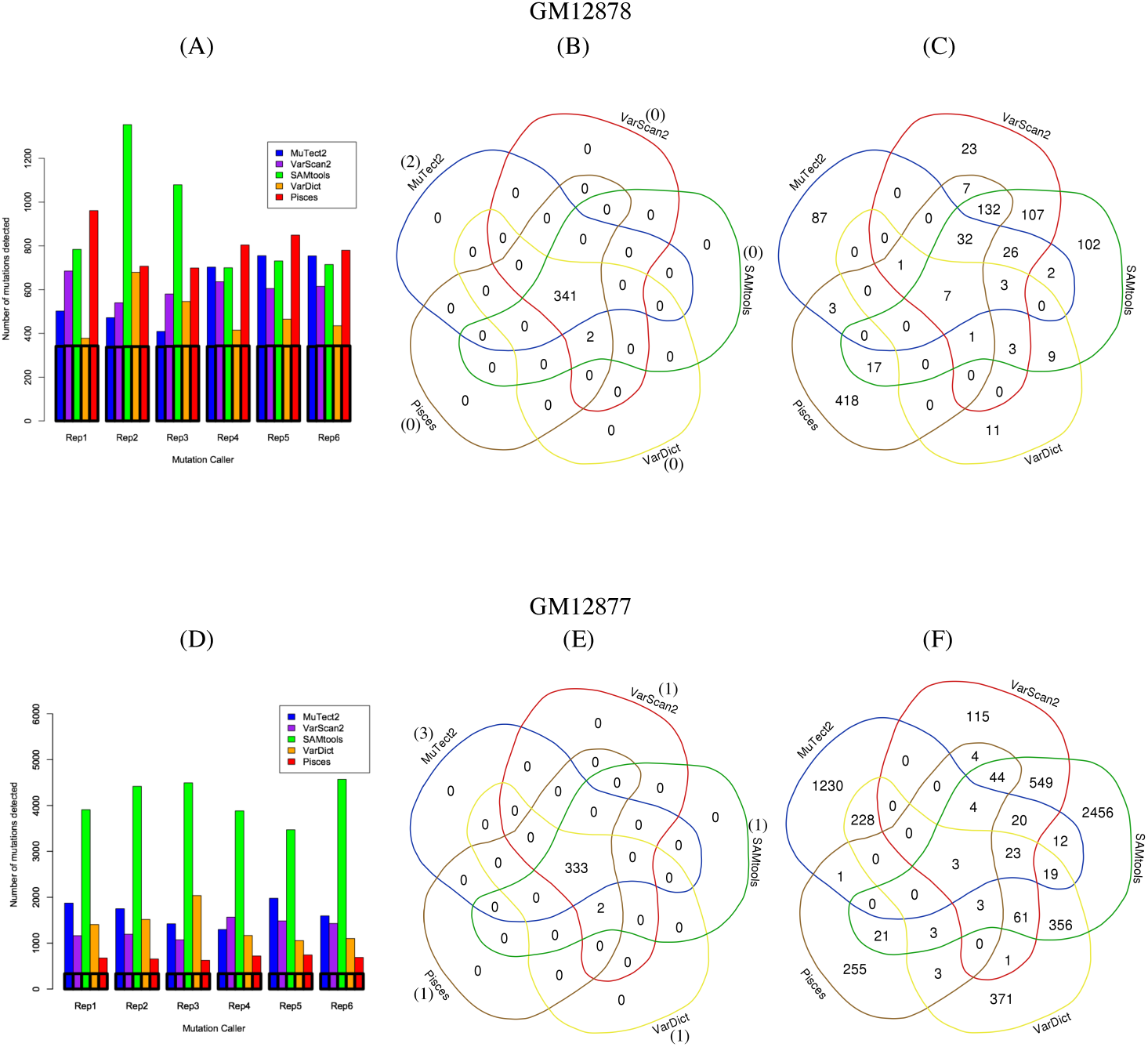
Comparison of five SNV calling approaches on TST 170 data sets using harmonized parameters. (A) Numbers of variants called by each algorithm, (B,C) distribution of true positive and false positives detections by the different algorithms. The numbers in parentheses represent total false negatives for each variant caller, i.e., the number of true variants not detected by the corresponding algorithm. (D-F)Same for GM12877.

We see similar overall trends in the Oncomine Focus data, as shown in Figure 6. SAMtools calls fewer variants after harmonization, while VarDict calls more. Most gold-standard variants are still identified, while false positives are greatly reduced. Still, the numbers of false positives are rather higher than would be desired, particularly for clinical sequencing where accuracy is key.

**Figure 6:**
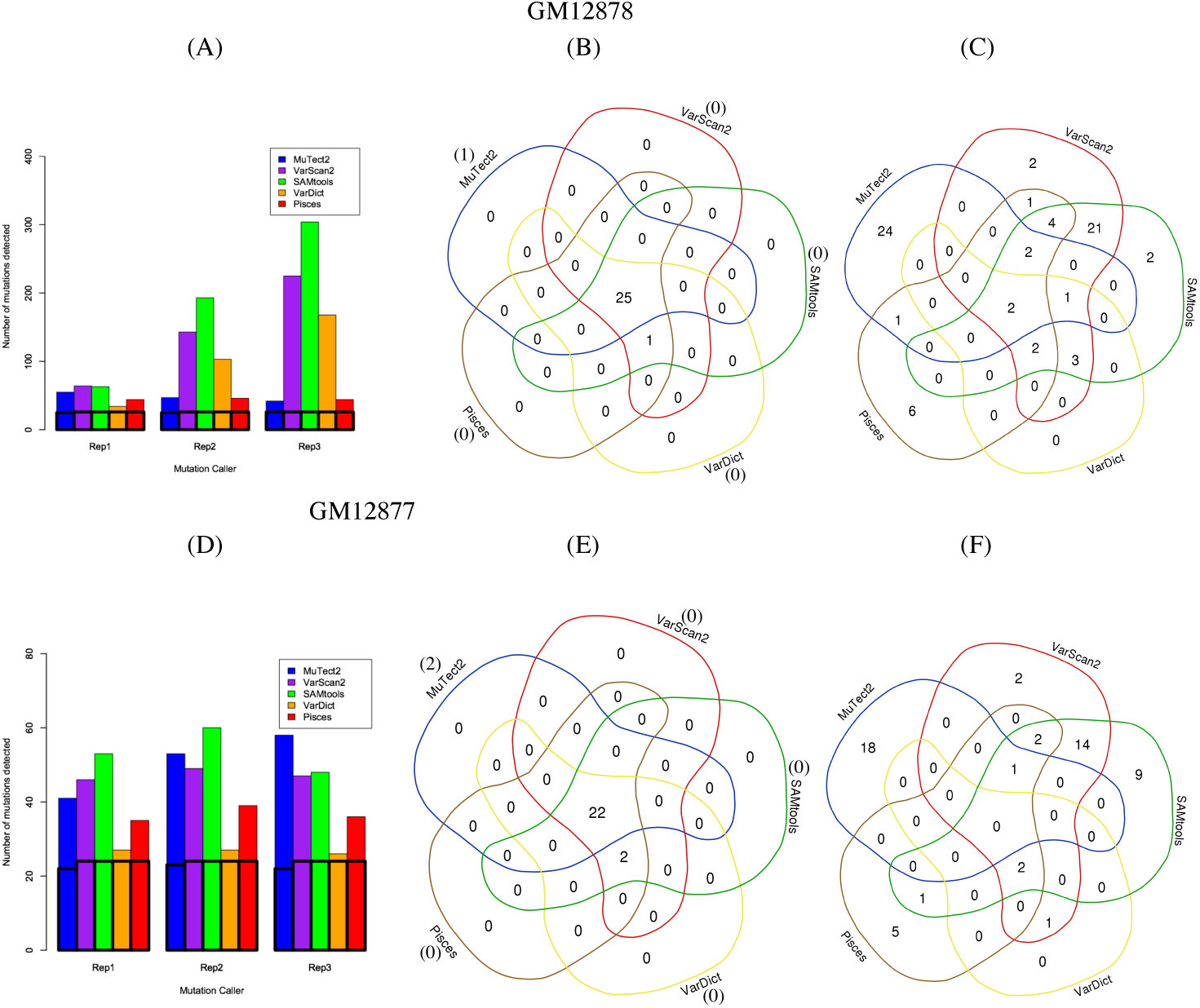
Comparison of five SNV calling approaches on Oncomine data sets using harmonized parameters. (A) Numbers of variants called by each algorithm, (B,C) distribution of true positive and false positives detections by the different algorithms. The numbers in parentheses represent total false negatives for each variant caller, i.e., the number of true variants not detected by the corresponding algorithm. (D-F) Same for GM12877.

Tables 2 to 5 report the complete true positive, false positive, false negative, precision and sensitivity numbers for each algorithm on each dataset, while Figure 7 summarizes precision and sensitivity visually. In terms of sensitivity and precision, there is no absolute winner. In two conditions, the TruSight170 GM12877 data (panel B) and the Oncomine Focus GM12878 data (panel C), Pisces is tied for first in sensitivity while offering higher precision. For the Oncomine Focus GM12877 data (panel D), it is VarDict that offers the highest precision while still maintaining perfect sensitivity. For the TruSight 170 GM12878 data (panel A), VarDict has the highest average precision but falls slightly short of the sensitivity of Pisces and SAMtools, although these have much lower precision. VarScan2 has the same (Oncomine) or slightly lower (TruSight 170) sensitivity as Pisces, VarDict and SAMtools, with intermediate precision. MuTect2 misses slightly more gold-standard SNV than the other algorithms, so ends up with the lowest sensitivity. It has variable precision.

**Table 2:** Variant calling performance with harmonized parameters on TST170-GM12878, in terms of true positives (TP), false positives (FP), false negatives (FN), precision (TP/(TP+FP)), and sensitivity (TP/(TP+FN)). Reported precision and sensitivity are averaged over the replicates.

|  | Rep 1<br>TP,FP,FN | Rep 2<br>TP,FP,FN | Rep 3<br>TP,FP,FN | Rep 4<br>TP,FP,FN | Rep 5<br>TP,FP,FN | Rep 6<br>TP,FP,FN | Prec | Sens |
| --- | --- | --- | --- | --- | --- | --- | --- | --- |
| MuTect2 | 341 161 2 | 337 135 6 | 339 70 4 | 340 363 3 | 340 415 3 | 340 414 3 | 0.601 | 0.990 |
| VarScan2 | 343 342 0 | 338 202 5 | 341 239 2 | 343 293 0 | 343 262 0 | 343 272 0 | 0.563 | 0.997 |
| SAMtools | 343 441 0 | 340 1014 3 | 342 737 1 | 343 357 0 | 343 388 0 | 343 372 0 | 0.407 | 0.998 |
| VarDict | 343 35 0 | 340 339 3 | 342 204 1 | 343 72 0 | 342 123 1 | 343 92 0 | 0.731 | 0.998 |
| Pisces | 343 618 0 | 340 367 3 | 342 357 1 | 343 461 0 | 343 506 0 | 343 437 0 | 0.433 | 0.998 |

**Table 3:**
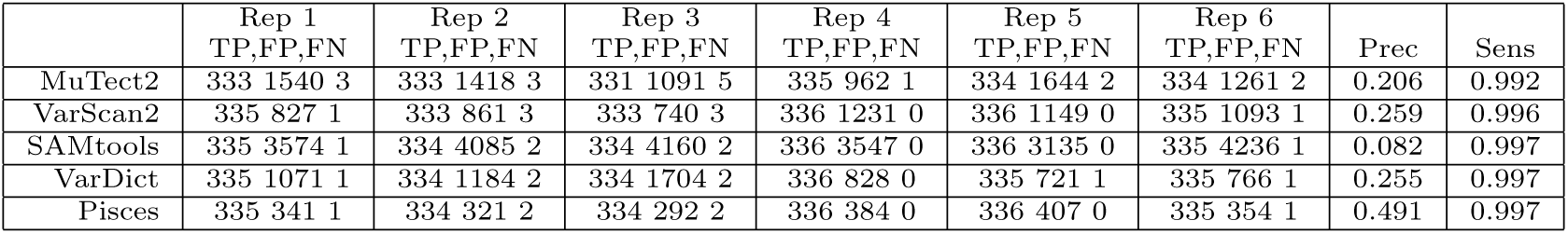
Variant calling performance with harmonized parameters on TST170-GM12877.

|  | Rep 1<br>TP,FP,FN | Rep 2<br>TP,FP,FN | Rep 3<br>TP,FP,FN | Rep 4<br>TP,FP,FN | Rep 5<br>TP,FP,FN | Rep 6<br>TP,FP,FN | Prec | Sens |
| --- | --- | --- | --- | --- | --- | --- | --- | --- |
| MuTect2 | 333 1540 3 | 333 1418 3 | 331 1091 5 | 335 962 1 | 334 1644 2 | 334 1261 2 | 0.206 | 0.992 |
| VarScan2 | 335 827 1 | 333 861 3 | 333 740 3 | 336 1231 0 | 336 1149 0 | 335 1093 1 | 0.259 | 0.996 |
| SAMtools | 335 3574 1 | 334 4085 2 | 334 4160 2 | 336 3547 0 | 336 3135 0 | 335 4236 1 | 0.082 | 0.997 |
| VarDict | 335 1071 1 | 334 1184 2 | 334 1704 2 | 336 828 0 | 335 721 1 | 335 766 1 | 0.255 | 0.997 |
| Pisces | 335 341 1 | 334 321 2 | 334 292 2 | 336 384 0 | 336 407 0 | 335 354 1 | 0.491 | 0.997 |

**Table 4:** Variant calling performance with harmonized parameters on Oncomine-GM12878.

|  | Rep 1<br>TP,FP,FN | Rep 2<br>TP,FP,FN | Rep 3<br>TP,FP,FN | Prec | Sens |
| --- | --- | --- | --- | --- | --- |
| MuTect2 | 25 30 1 | 26 21 0 | 26 16 0 | 0.542 | 0.987 |
| VarScan2 | 26 38 0 | 26 117 0 | 26 199 0 | 0.235 | 1.000 |
| SAMtools | 26 37 0 | 26 167 0 | 26 278 0 | 0.211 | 1.000 |
| VarDict | 26 8 0 | 26 77 0 | 26 142 0 | 0.391 | 1.000 |
| Pisces | 26 18 0 | 26 20 0 | 26 18 0 | 0.582 | 1.000 |

**Table 5:** Variant calling performance with harmonized parameters on Oncomine-GM12877.

|  | Rep 1<br>TP,FP,FN | Rep 2<br>TP,FP,FN | Rep 3<br>TP,FP,FN | Prec | Sens |
| --- | --- | --- | --- | --- | --- |
| MuTect2 | 22 19 2 | 23 30 1 | 22 36 2 | 0.450 | 0.931 |
| VarScan2 | 24 22 0 | 24 25 0 | 24 23 0 | 0.507 | 1.000 |
| SAMtools | 24 29 0 | 24 36 0 | 24 24 0 | 0.451 | 1.000 |
| VarDict | 24 3 0 | 24 3 0 | 24 2 0 | 0.900 | 1.000 |
| Pisces | 24 11 0 | 24 15 0 | 24 12 0 | 0.656 | 1.000 |

**Figure 7:**
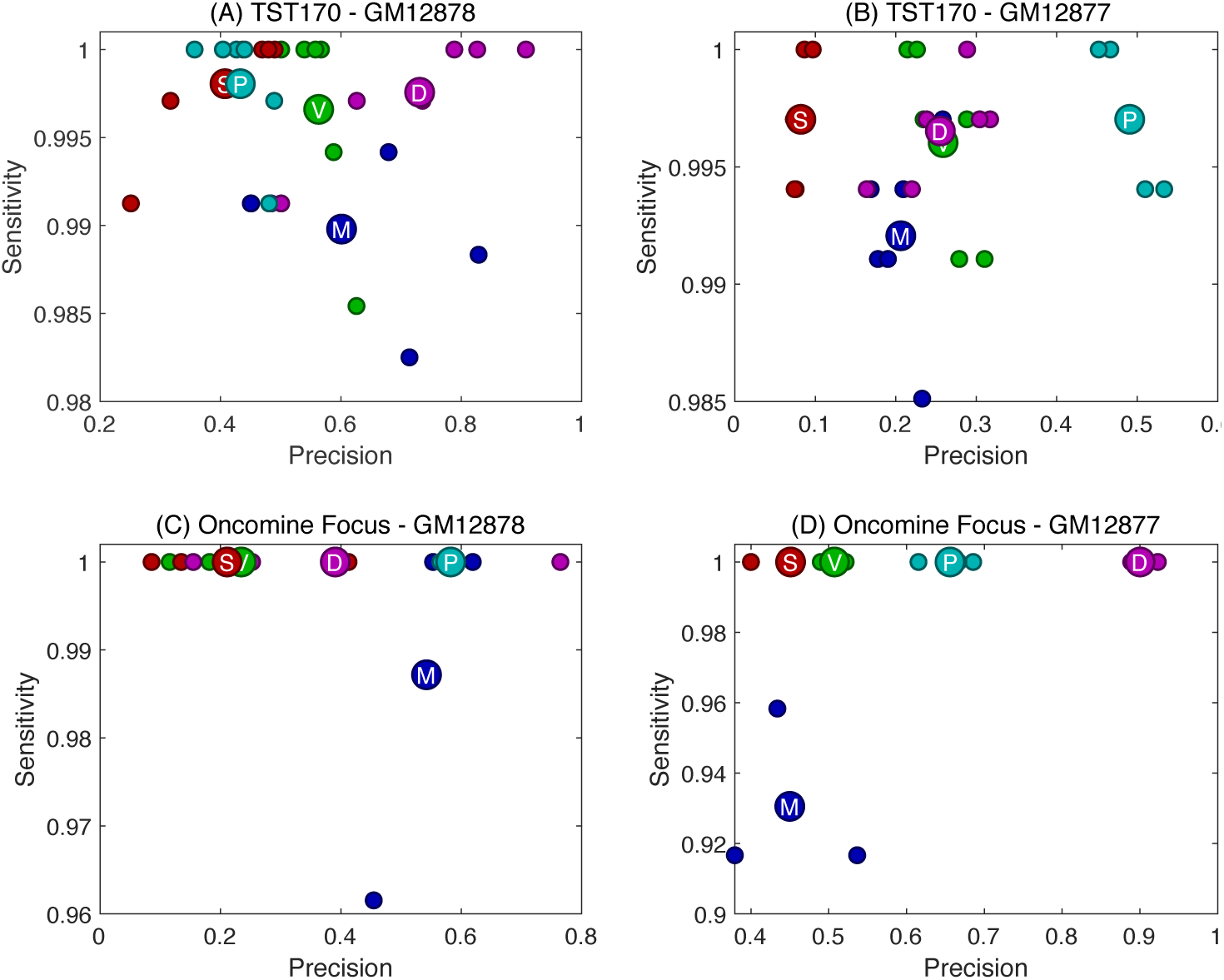
Precision and Sensitivity of the five variant callers: MuTect2 (M), SAMtools (S), VarScan2 (V), Pisces (P) and VarDict (D). Larger circles represent mean precision and sensitivity across replicates, while smaller circles represent performance on individual replicates.

### Thresholding depth and allele frequency is inadequate for removing false positives

To investigate further the false positive variant calls, we focused on replicate 1 of each of the four conditions (GM12878 and GM12877, sequencing using TST170 or Oncomine Focus panels). Figure 8 shows the false positives called by each algorithm as a function of read coverage and allele frequency. Unsurprisingly, false positives tend to come at lower coverages and/or allele frequencies. However, there is a substantial number of errors at, for example, coverage of 500 or more reads and allele frequency of 0.1 or more, which would normally be considered well-supported. We also see that, for the most part, the errors of the algorithms are intermingled. One algorithm will make an error at a given coverage and allele frequency, while another algorithm avoids that error. Yet the second algorithm will make a different error at a site with similar coverage and allele frequency. Thus, other features besides these two are driving differences in the errors between algorithms. We also see a very small number of sites with high allele frequencies, above 0.4 say, and high coverage, 500 or 1000 reads or more. Again, however, only some algorithms call these variants. From these plots, and comparing to Figure 1, it is clear that stricter thresholding on coverage and/or allele frequency is not a sufficient means to eliminate false positives or eliminate algorithm disagreements. For example, from Figure 8A we observe that to eliminate all false positives we could choose a coverage threshold of 1500 and a minimum allele frequency of 0.25, selecting out the empty upper right corner of the plot. However, from Figure 1A, we see that applying the same thresholds would eliminate from consideration a large number of the gold-standard mutations, resulting in an unacceptable loss of true positives.

**Figure 8:**
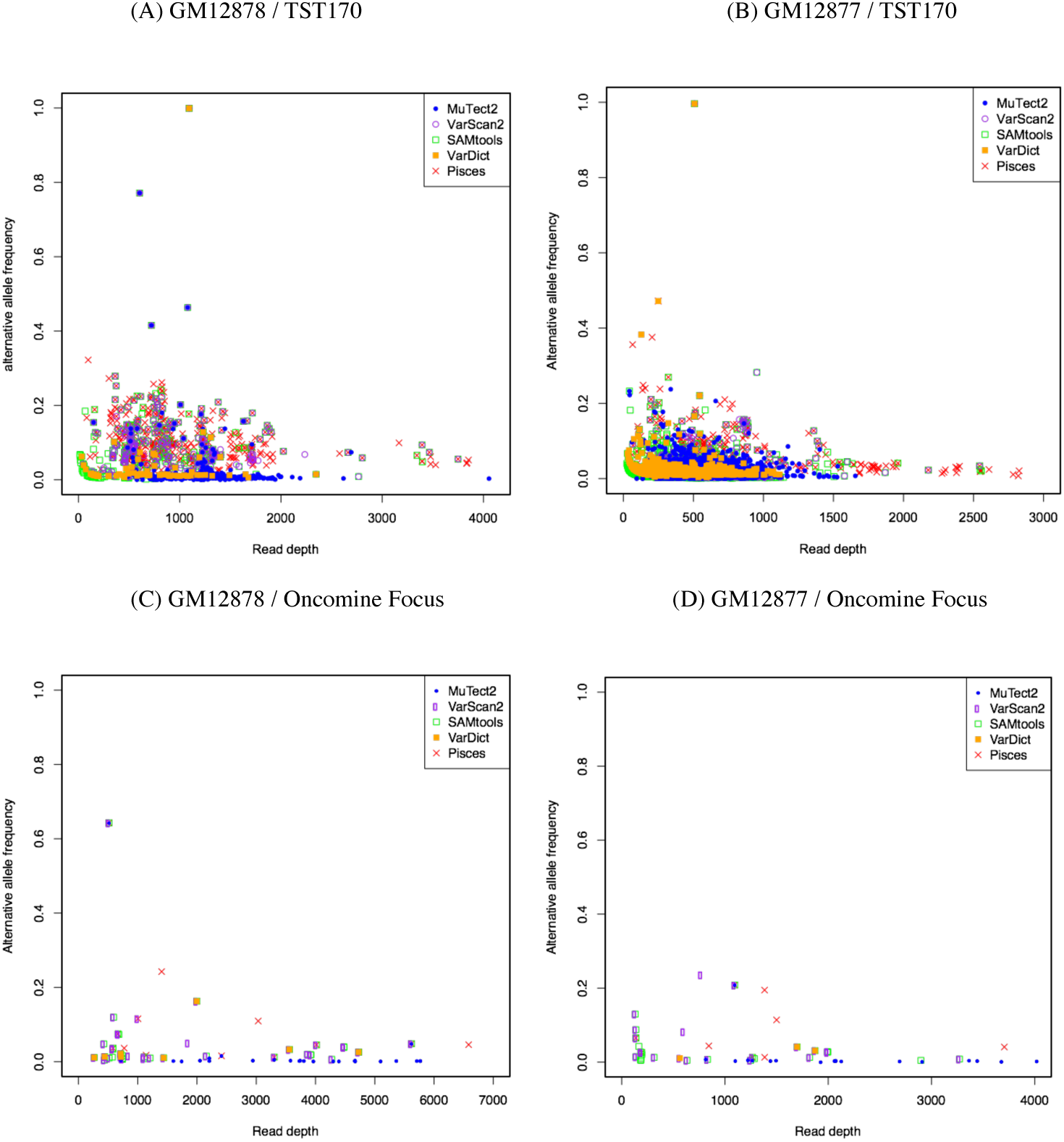
False positive variant calls by each algorithm, as a function of coverage and allele frequency, for replicate 1 of each condition.

To formalize the trade offs involved in thresholding on depth and allele frequency, we post-filtered all variant calls at 10 different levels of stringency: a minimum of 20 reads and allele frequency 0.01, 40 reads and allele frequency 0.02, 60 reads and allele frequency 0.03, and so on up to requiring a minimum of 200 reads and allele frequency 0.10. The least stringent threshold is almost equivalent to the results reported above, as our harmonized variant calling parameters already included a minimum allele frequency of 0.01, and very few mutations can be called with fewer than 20 reads. The most stringent threshold would be considered unreasonably stringent by many practitioners—firstly, because mutations with allele frequency less than 0.10 can be clinical relevant, and secondly because there is a general consensus that even baseline clinical sequencing approaches ought to be able to detect mutations down to at least 0.05 allele frequency. Figure 9 shows the number of true positives and false positives, averaged across replicates, obtained at these different levels of stringency. For example, consider the performance of SAMtools on the GM12878 TST170 data in panel A. Increasing depth and allele frequency stringency can reduce false positives from an average of over 400 per replicate to near 100—a substantial improvement. This comes at the loss, however, of roughly 15 true positives per replicate, and still leaves us with a substantial empirical false discovery rate of approximately 23%. Results are better for some algorithms and worse for others, and results vary depending on the dataset. Only for the GM12877 Oncomine Focus data do we see that increasing stringency eliminates false positives with no loss of true positives. However, it is important to recall that the true allele frequencies of all mutations in our cell line data are at least 0.5, and the empirical allele frequencies of gold standard mutations in the Oncomine Focus data sets never fell below 0.4. In tumor data, where there is typically a broader range of allele frequencies, including nearer to zero, one would expect thresholding on depth and allele frequency to be less helpful than we see in Figure 9.

**Figure 9:**
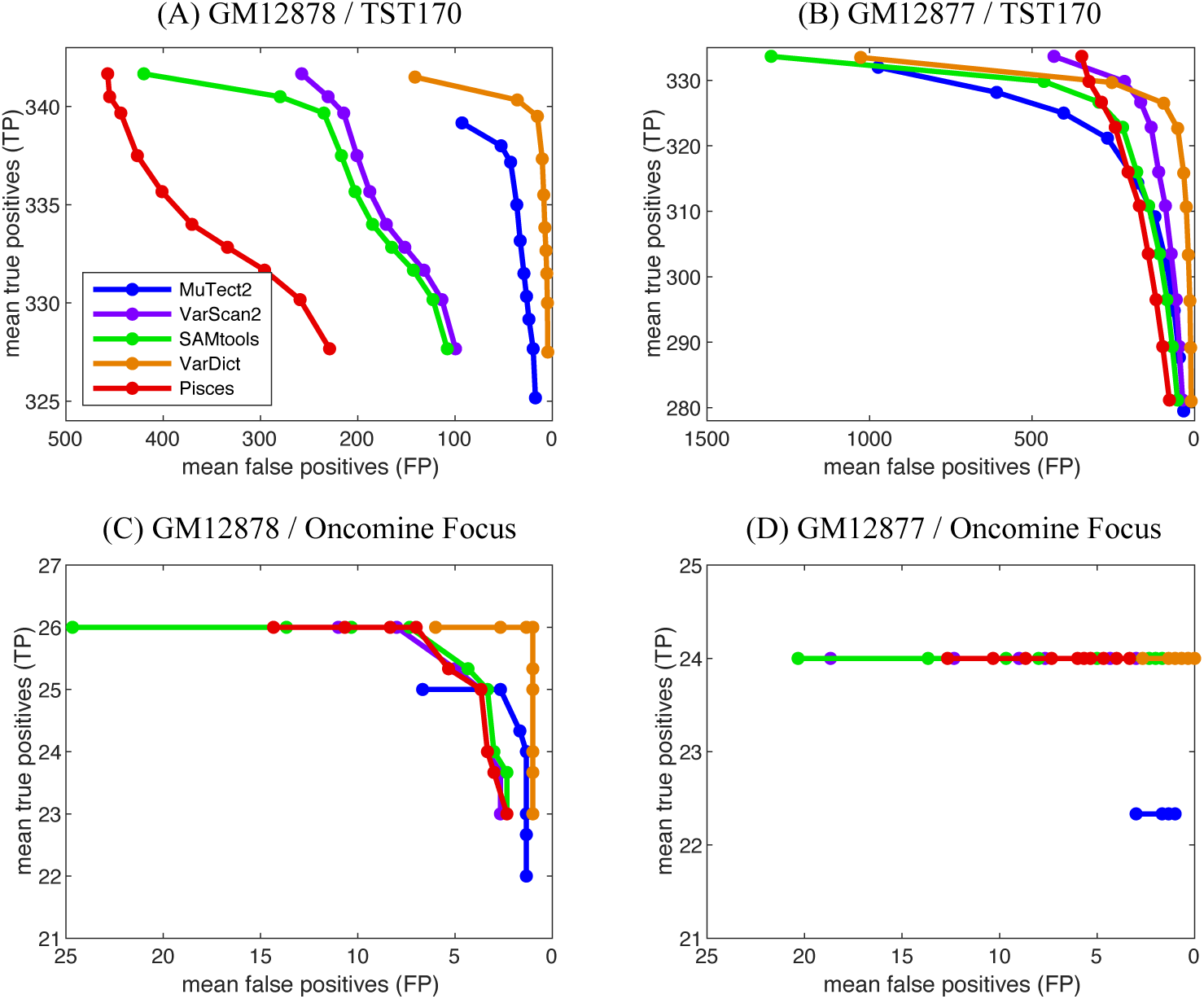
True and false positive calls by each algorithm and for each condition (GM12878 or GM12877, assayed with TST170 or Oncomine Focus panels), average over replicates, when calling stringency is increased by ranging minimum read depth from 20 to 200 simultaneously with allele frequency ranging from 0.01 to 0.10.

### Intersecting mutation callers’ results reduces false positives, while maintaining sensitivity

From Figures 5 and 6, it is clear that each individual mutation caller has reported many false positives. Intersecting at least two mutation callers can remarkably reduce the number of such false positives. For example, the list of known mutations created by [17] applies the same idea to the calls made by FreeBayes, Platypus GATK3 and Strelka. To take an example from our data, in Figure 5C we observe that in replicate 1 of TST170-GM12878, VarDict makes 26 false positive calls that are not made by Pisces, and Pisces makes 609 false positive calls that are not made by VarDict. At the same time, from the corresponding Venn diagram in Figure 5B, it is evident that both callers are able to detect the 343 known mutations. Thus, intersecting these two mutation callers can exclude a significant percentage of the false positives. In fact, it reduces the number of false positives to 9, and an EFDR of just 2.56%. This raises questions such as how many mutation callers should be combined or what are the best combinations of mutation callers in order to get best performance.

We analyzed all the data sets and calculated the performance obtained by intersecting every possible combination of the five programs’ SNV calls. Tables 6 to 9 present the results in detail. From the tables, we observe that intersecting exactly two mutation callers leads to an improved performance. For example, for replicate 1 of TST170-GM12878 data, intersecting MuTect2 and VarDict leads to 341 true positives and only 11 false positives (with an EFDR of 3%). It seems that interesting MuTect2 and VarDict leads to the lowest false detection but intersecting VarScan2 and SAMtools, or SAMtools and Pisces, MuTect2 and VarScan2, or MuTect2 and SAMtools leads to many false positives. This suggests some kind of complementarity in the errors made by MuTect2 and VarDict, that is not shared by other algorithm combinations. Table 6 also shows that intersecting three of the five mutation callers leads to an improved performance. For example, for replicate 1 of TST170-GM12878 data, combining VarDict, Pisces and SAMtools leads to 343 true detections and 8 false positives (with an EFDR of 2.2%). Intersecting VarDict, Pisces and MuTect2 leads to 341 true detections and 8 false positives (with an EFDR of 2.3%). Analyzing the number of true and false detections in any combination of the mutation callers across all replicates suggests that the combination of VarDict, Pisces and MuTect2 (if the interest is in choosing 3 out 5 callers) or the combination of VarDict and Pisces (if the interest is in choosing 2 out 5 callers) significantly improves the performance compared to each individual caller.

**Table 6:**
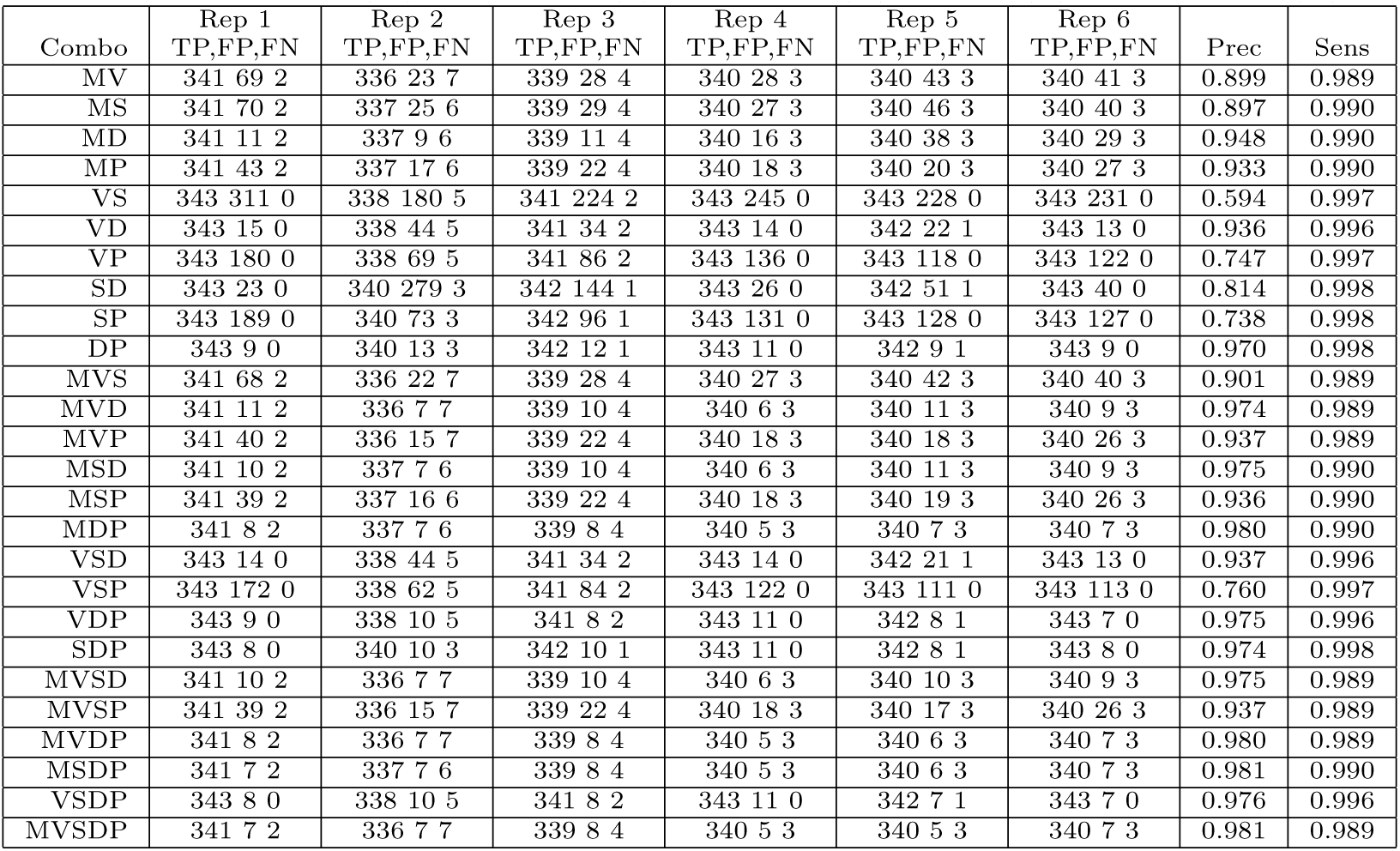
Performance of algorithm combinations on TST170-GM12878 data. Algorithms are coded as: MuTect2 (M), SAMtools (S), VarScan2 (V), Pisces (P) and VarDict (D).

**Table 7:** Performance of algorithm combinations on TST170-GM12877 data. Algorithms are coded as: MuTect2 (M), SAMtools (S), VarScan2 (V), Pisces (P) and VarDict (D).

| Combo | Rep 1<br>TP,FP,FN | Rep 2<br>TP,FP,FN | Rep 3<br>TP,FP,FN | Rep 4<br>TP,FP,FN | Rep 5<br>TP,FP,FN | Rep 6<br>TP,FP,FN | Prec | Sens |
| --- | --- | --- | --- | --- | --- | --- | --- | --- |
| MV | 333 50 3 | 332 71 4 | 330 60 6 | 335 57 1 | 334 83 2 | 334 60 2 | 0.840 | 0.991 |
| MS | 333 81 3 | 333 90 3 | 331 75 5 | 335 74 1 | 334 98 2 | 334 79 2 | 0.801 | 0.992 |
| MD | 333 273 3 | 333 295 3 | 331 303 5 | 335 206 1 | 334 240 2 | 334 225 2 | 0.567 | 0.992 |
| MP | 333 8 3 | 333 12 3 | 331 6 5 | 335 12 1 | 334 18 2 | 334 16 2 | 0.965 | 0.992 |
| VS | 335 707 1 | 333 732 3 | 333 624 3 | 336 920 0 | 336 870 0 | 335 902 1 | 0.300 | 0.996 |
| VD | 335 91 1 | 333 117 3 | 333 191 3 | 336 110 0 | 335 83 1 | 335 87 1 | 0.752 | 0.996 |
| VP | 335 58 1 | 333 67 3 | 333 58 3 | 336 87 0 | 336 101 0 | 335 87 1 | 0.816 | 0.996 |
| SD | 335 468 1 | 334 572 2 | 334 962 2 | 336 347 0 | 335 223 1 | 335 300 1 | 0.444 | 0.997 |
| SP | 335 78 1 | 334 88 2 | 334 73 2 | 336 114 0 | 336 113 0 | 335 116 1 | 0.777 | 0.997 |
| DP | 335 12 1 | 334 14 2 | 334 15 2 | 336 13 0 | 335 18 1 | 335 13 1 | 0.959 | 0.997 |
| MVS | 333 50 3 | 332 70 4 | 330 59 6 | 335 55 1 | 334 79 2 | 334 58 2 | 0.844 | 0.991 |
| MVD | 333 26 3 | 332 43 4 | 330 41 6 | 335 29 1 | 334 38 2 | 334 31 2 | 0.906 | 0.991 |
| MVP | 333 7 3 | 332 10 4 | 330 6 6 | 335 8 1 | 334 16 2 | 334 13 2 | 0.971 | 0.991 |
| MSD | 333 45 3 | 333 52 3 | 331 50 5 | 335 36 1 | 334 44 2 | 334 36 2 | 0.884 | 0.992 |
| MSP | 333 7 3 | 333 10 3 | 331 6 5 | 335 8 1 | 334 16 2 | 334 13 2 | 0.971 | 0.992 |
| MDP | 333 3 3 | 333 4 3 | 331 4 5 | 335 3 1 | 334 10 2 | 334 7 2 | 0.985 | 0.992 |
| VSD | 335 90 1 | 333 117 3 | 333 190 3 | 336 109 0 | 335 83 1 | 335 84 1 | 0.753 | 0.996 |
| VSP | 335 54 1 | 333 65 3 | 333 55 3 | 336 79 0 | 336 94 0 | 335 83 1 | 0.825 | 0.996 |
| VDP | 335 6 1 | 333 10 3 | 333 9 3 | 336 9 0 | 335 14 1 | 335 11 1 | 0.971 | 0.996 |
| SDP | 335 9 1 | 334 12 2 | 334 12 2 | 336 11 0 | 335 16 1 | 335 12 1 | 0.965 | 0.997 |
| MVSD | 333 26 3 | 332 43 4 | 330 40 6 | 335 28 1 | 334 38 2 | 334 30 2 | 0.907 | 0.991 |
| MVSP | 333 7 3 | 332 10 4 | 330 6 6 | 335 8 1 | 334 16 2 | 334 13 2 | 0.971 | 0.991 |
| MVDP | 333 3 3 | 333 4 3 | 331 4 5 | 335 3 1 | 334 10 2 | 334 6 2 | 0.985 | 0.992 |
| MSDP | 333 3 3 | 332 4 4 | 330 4 6 | 335 3 1 | 334 10 2 | 334 6 2 | 0.985 | 0.991 |
| VSDP | 335 6 1 | 333 10 3 | 333 9 3 | 336 9 0 | 335 14 1 | 335 11 1 | 0.971 | 0.996 |
| MVSDP | 333 3 3 | 332 4 4 | 330 4 6 | 335 3 1 | 334 10 2 | 334 6 2 | 0.985 | 0.991 |

**Table 8:** Performance of algorithm combinations on Oncomine-GM12878 data. Algorithms are coded as: MuTect2 (M), SAMtools (S), VarScan2 (V), Pisces (P) and VarDict (D).

| Combo | Rep 1<br>TP,FP,FN | Rep 2<br>TP,FP,FN | Rep 3<br>TP,FP,FN | Prec | Sens |
| --- | --- | --- | --- | --- | --- |
| MV | 25 5 1 | 26 11 0 | 26 7 0 | 0.775 | 0.987 |
| MS | 25 5 1 | 26 11 0 | 26 8 0 | 0.767 | 0.987 |
| MD | 25 3 1 | 26 6 0 | 26 3 0 | 0.867 | 0.987 |
| MP | 25 5 1 | 26 3 0 | 26 2 0 | 0.886 | 0.987 |
| VS | 26 35 0 | 26 115 0 | 26 174 0 | 0.247 | 1.000 |
| VD | 26 8 0 | 26 67 0 | 26 124 0 | 0.406 | 1.000 |
| VP | 26 11 0 | 26 11 0 | 26 12 0 | 0.697 | 1.000 |
| SD | 26 8 0 | 26 67 0 | 26 107 0 | 0.413 | 1.000 |
| SP | 26 10 0 | 26 12 0 | 26 11 0 | 0.703 | 1.000 |
| DP | 26 4 0 | 26 3 0 | 26 3 0 | 0.887 | 1.000 |
| MVS | 25 5 1 | 26 11 0 | 26 7 0 | 0.775 | 0.987 |
| MVD | 25 3 1 | 26 6 0 | 26 3 0 | 0.867 | 0.987 |
| MVP | 25 4 1 | 26 3 0 | 26 2 0 | 0.896 | 0.987 |
| MSD | 25 3 1 | 26 6 0 | 26 3 0 | 0.867 | 0.987 |
| MSP | 25 4 1 | 26 3 0 | 26 2 0 | 0.896 | 0.987 |
| MDP | 25 2 1 | 26 2 0 | 26 1 0 | 0.939 | 0.987 |
| VSD | 26 8 0 | 26 65 0 | 26 106 0 | 0.416 | 1.000 |
| VSP | 26 10 0 | 26 11 0 | 26 11 0 | 0.709 | 1.000 |
| VDP | 26 4 0 | 26 2 0 | 26 2 0 | 0.908 | 1.000 |
| SDP | 26 4 0 | 26 2 0 | 26 2 0 | 0.908 | 1.000 |
| MVSD | 25 3 1 | 26 6 0 | 26 3 0 | 0.867 | 0.987 |
| MVSP | 25 4 1 | 26 3 0 | 26 2 0 | 0.896 | 0.987 |
| MVDP | 25 2 1 | 26 2 0 | 26 1 0 | 0.939 | 0.987 |
| MSDP | 25 2 1 | 26 2 0 | 26 1 0 | 0.939 | 0.987 |
| VSDP | 26 4 0 | 26 2 0 | 26 2 0 | 0.908 | 1.000 |
| MVSDP | 25 2 1 | 26 2 0 | 26 1 0 | 0.939 | 0.987 |

**Table 9:** Performance of algorithm combinations on Oncomine-GM12877 data. Algorithms are coded as: MuTect2 (M), SAMtools (S), VarScan2 (V), Pisces (P) and VarDict (D).

| Combo | Rep 1<br>TP,FP,FN | Rep 2<br>TP,FP,FN | Rep 3<br>TP,FP,FN | Prec | Sens |
| --- | --- | --- | --- | --- | --- |
| MV | 22 1 2 | 23 4 1 | 22 3 2 | 0.896 | 0.931 |
| MS | 22 1 2 | 23 4 1 | 22 3 2 | 0.896 | 0.931 |
| MD | 22 0 2 | 23 2 1 | 22 1 2 | 0.959 | 0.931 |
| MP | 22 1 2 | 23 5 1 | 22 3 2 | 0.886 | 0.931 |
| VS | 24 19 0 | 24 24 0 | 24 21 0 | 0.530 | 1.000 |
| VD | 24 3 0 | 24 2 0 | 24 2 0 | 0.912 | 1.000 |
| VP | 24 5 0 | 24 7 0 | 24 8 0 | 0.784 | 1.000 |
| SD | 24 2 0 | 24 2 0 | 24 2 0 | 0.923 | 1.000 |
| SP | 24 6 0 | 24 7 0 | 24 7 0 | 0.783 | 1.000 |
| DP | 24 2 0 | 24 2 0 | 24 1 0 | 0.935 | 1.000 |
| MVS | 22 1 2 | 23 4 1 | 22 3 2 | 0.896 | 0.931 |
| MVD | 22 0 2 | 23 2 1 | 22 1 2 | 0.959 | 0.931 |
| MVP | 22 1 2 | 23 4 1 | 22 3 2 | 0.896 | 0.931 |
| MSD | 22 0 2 | 23 2 1 | 22 1 2 | 0.959 | 0.931 |
| MSP | 22 1 2 | 23 4 1 | 22 3 2 | 0.896 | 0.931 |
| MDP | 22 0 2 | 23 2 1 | 22 1 2 | 0.959 | 0.931 |
| VSD | 24 2 0 | 24 2 0 | 24 2 0 | 0.923 | 1.000 |
| VSP | 24 5 0 | 24 6 0 | 24 7 0 | 0.801 | 1.000 |
| VDP | 24 2 0 | 24 2 0 | 24 1 0 | 0.935 | 1.000 |
| SDP | 24 2 0 | 24 2 0 | 24 1 0 | 0.935 | 1.000 |
| MVSD | 22 0 2 | 23 2 1 | 22 1 2 | 0.959 | 0.931 |
| MVSP | 22 1 2 | 23 4 1 | 22 3 2 | 0.896 | 0.931 |
| MVDP | 22 0 2 | 23 2 1 | 22 1 2 | 0.959 | 0.931 |
| MSDP | 22 0 2 | 23 2 1 | 22 1 2 | 0.959 | 0.931 |
| VSDP | 24 2 0 | 24 2 0 | 24 1 0 | 0.935 | 1.000 |
| MVSDP | 22 0 2 | 23 2 1 | 22 1 2 | 0.959 | 0.931 |

Figure 10 summarizes mean precision and sensitivity for select combinations. Intersecting the results of two or more algorithms cannot increase the number of true positive detections; the number can only stay the same or decrease. Thus, the sensitivity of a combination is no higher than that of its individual algorithms. By ruling out false positives, however, precision can increase. So a combination is good to the extent that it remains at high sensitivity while increasing precision. We see, for example, intersecting SAMtools and Pisces (SP) loses no sensitivity on these datasets, while substantially improving precision. Still, that precision remains between 0.7 and 0.8. The VarDict-Pisces combination (DP) has equal or slightly lower sensitivity (due to VarDict), but much better precision, at over 0.9. Indeed this was the highest-precision pair for all four conditions. We can intersect even more algorithms, including all five, to obtain yet higher precision, but only slightly, and at substantial cost of sensitivity due to the inclusion of MuTect2.

**Figure 10:**
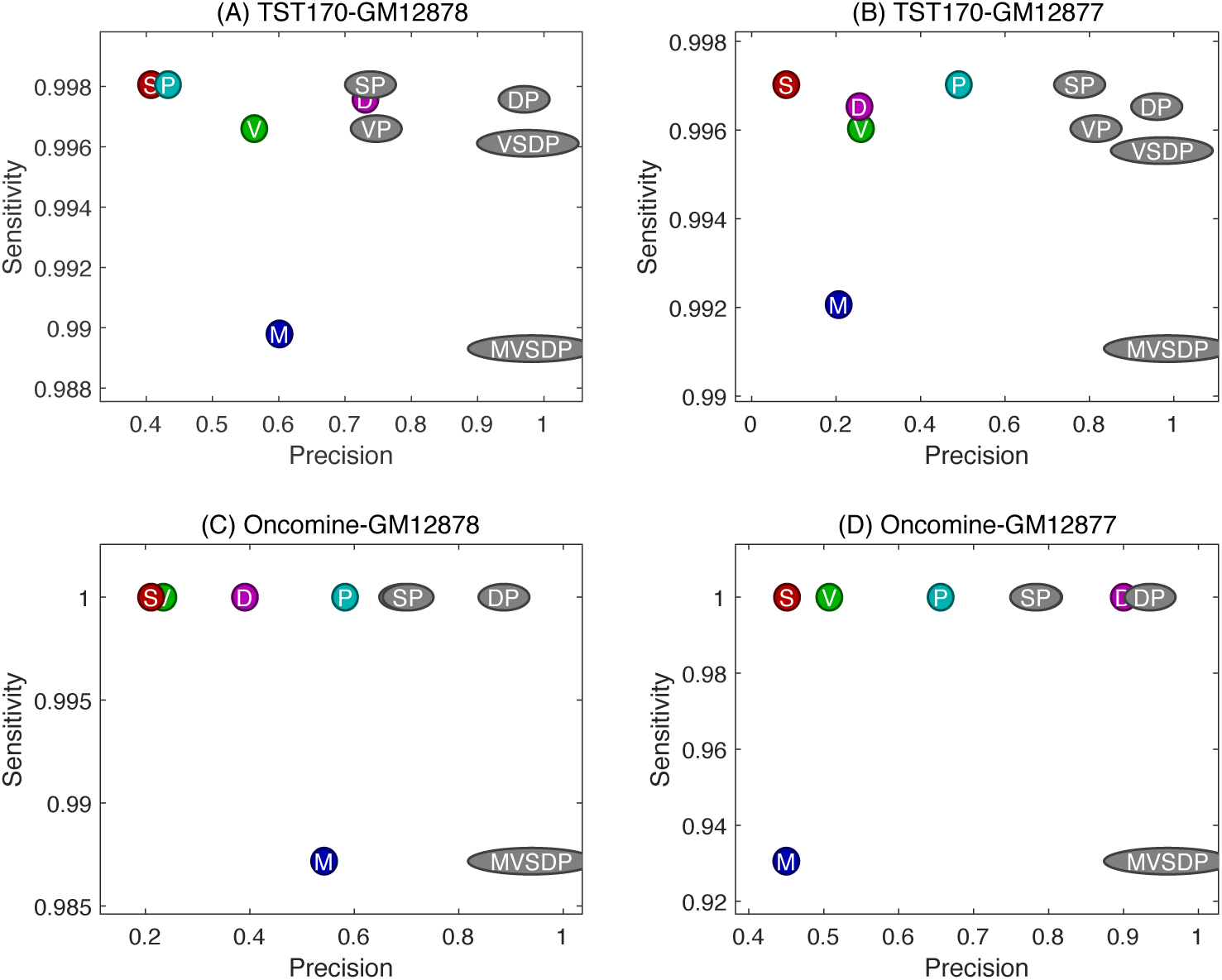
Mean precision and sensitivity values by individual variant callers and some selected combinations. M, V, S, D and P stands for MuTect2, VarScan2, SAMtools, VarDict and Pisces, respectively. Colored circles refer to individual algorithms, while grey ellipses refer to intersections of results from the specified algorithms.

### Replicate analysis increases the accuracy of an individual mutation caller

We also analyzed the data by combining the mutations detected by an individual caller across the replicates. Tables 10 to 13 report true positives, false positives, false negatives, precision and sensitivity if variant calls are only accepted when they appear in at least *N* replicates. For example, from Table 10, we observe that for TST170-GM12878 data, when we looked at common mutations detected by MuTect2 in at least two of six replicates, we get 341 TPs, 82 FPs, and 2 FNs. By comparison, the average performance across single replicates was 339.5 TPs, 259.6 FPs, and 3.5 FNs.

**Table 10:** Performance of intersecting two or more replicates to obtain variant calls on the TST170-GM12878 data.

| | $\geq 2$ reps<br>TP,FP,FN<br>Prec,Sens | $\geq 3$ reps<br>TP,FP,FN<br>Prec,Sens | $\geq 4$ reps<br>TP,FP,FN<br>Prec,Sens | $\geq 5$ reps<br>TP,FP,FN<br>Prec,Sens | all 6 reps<br>TP,FP,FN<br>Prec,Sens |
| --- | --- | --- | --- | --- | --- |
| MuTect2 | 341 82 2<br>0.806 0.994 | 341 47 2<br>0.879 0.994 | 341 31 2<br>0.917 0.994 | 338 22 5<br>0.939 0.985 | 335 12 8<br>0.965 0.977 |
| VarScan2 | 343 304 0<br>0.530 1.000 | 343 216 0<br>0.614 1.000 | 343 148 0<br>0.699 1.000 | 341 114 2<br>0.749 0.994 | 338 80 5<br>0.809 0.985 |
| SAMtools | 343 379 0<br>0.475 1.000 | 343 225 0<br>0.604 1.000 | 343 149 0<br>0.697 1.000 | 342 112 1<br>0.753 0.997 | 340 74 3<br>0.821 0.991 |
| VarDict | 343 24 0<br>0.935 1.000 | 343 11 0<br>0.969 1.000 | 343 8 0<br>0.977 1.000 | 342 7 1<br>0.980 0.997 | 339 6 4<br>0.983 0.988 |
| Pisces | 343 659 0<br>0.342 1.000 | 343 455 0<br>0.430 1.000 | 343 297 0<br>0.536 1.000 | 342 168 1<br>0.671 0.997 | 340 100 3<br>0.773 0.991 |

**Table 11:**
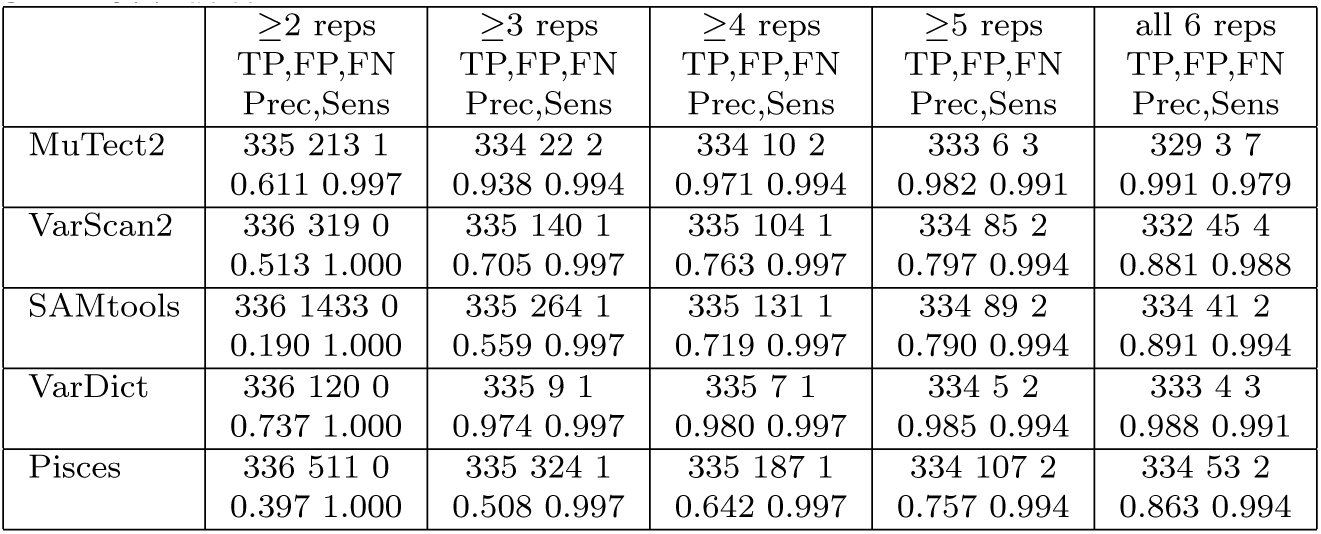
Performance of intersecting two or more replicates to obtain variant calls on the TST170-GM12877 data.

**Table 12:** Performance of intersecting two or more replicates to obtain variant calls on the Oncomine-GM12878 data.

| | $\geq 2$ reps<br>TP,FP,FN<br>Prec,Sens | all 3 reps<br>TP,FP,FN<br>Prec,Sens |
| --- | --- | --- |
| MuTect2 | 26 8 0<br>0.765 1.000 | 25 3 1<br>0.893 0.962 |
| VarScan2 | 26 34 0<br>0.433 1.000 | 26 7 0<br>0.788 1.000 |
| SAMtools | 26 48 0<br>0.351 1.000 | 26 11 0<br>0.703 1.000 |
| VarDict | 26 7 0<br>0.788 1.000 | 26 3 0<br>0.897 1.000 |
| Pisces | 26 13 0<br>0.667 1.000 | 26 3 0<br>0.897 1.000 |

**Table 13:** Performance of intersecting two or more replicates to obtain variant calls on the Oncomine-GM12877 data.

| | $\geq 2$ reps<br>TP,FP,FN<br>Prec,Sens | all 3 reps<br>TP,FP,FN<br>Prec,Sens |
| --- | --- | --- |
| MuTect2 | 23 8 1<br>0.742 0.958 | 21 4 3<br>0.840 0.875 |
| VarScan2 | 24 19 0<br>0.558 1.000 | 24 11 0<br>0.686 1.000 |
| SAMtools | 24 27 0<br>0.471 1.000 | 24 16 0<br>0.600 1.000 |
| VarDict | 24 2 0<br>0.923 1.000 | 24 1 0<br>0.960 1.000 |
| Pisces | 24 11 0<br>0.686 1.000 | 24 8 0<br>0.750 1.000 |

Figure 11 summarizes the precision and sensitivity. Requiring mutations to occur in increasing number of replicates *N* cannot improve sensitivity, but can improve precision. Generally, we see that requiring mutations to occur in more than one replicate increases precision without any cost to sensitivity up to some point, but that requiring mutations to occur in too many replicates (including all) can have a substantial sensitivity cost. On the other hand, we can push precision above 0.9, and on most datasets, near to 1.0. VarDict generally performs best in this scenario, occupying the upper right hand corner of the plots, although for the Oncomine-GM12878 dataset, the strict three-way intersection of Pisces results performs identically with the three-way VarDict intersection. (This cannot be seen from the plot, because the Pisces P3 bubble is on top of the VarDict D3 bubble, but the result is apparent from Table 12.) Also importantly, the best performances obtained by the replicate analysis exceed the performance of intersecting different variant callers results. For example, on the TST170-GM12878 data, the best algorithm intersection was VarDict-Pisces, with mean precision 0.935 and sensitivity 1.0. In the replicate analysis, focusing on VarDict results that appear in at least three replicates produces precision 0.977 and sensitivity 1.0.

**Figure 11:**
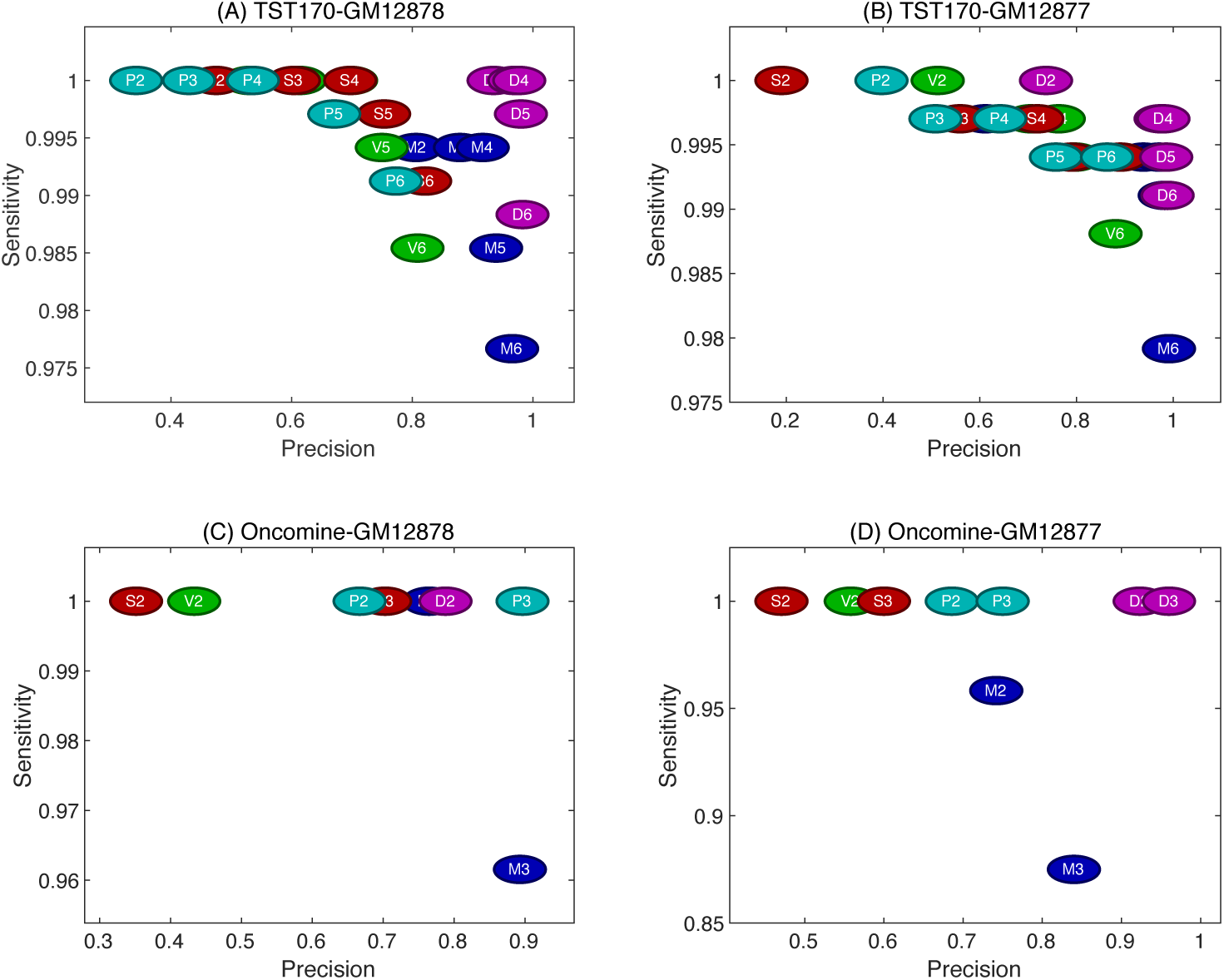
Precision and sensitivity when each variant caller’s results are filtered so that they must appear in at least 2 replicates, or at least 3, etc. Each bubble specifies an algorithm (M=MuTect2, V=VarScan2, S=SAMtools, D=VarDict, P=Pisces) and the minimum number of replicates in which a variant call appears.

## Conclusions

In this work, we looked at factors contributing to the success of SNV calling in clinical-style samples, including different sequencing platforms, targeting panels, variant callers, and replication. We generally found similar results for the TruSight 170 data sequenced on an Illumina platform and for the Oncomine Focus data sequenced on an Ion Torrent platform.

Thus, we concluded that sequencing platform and targeting panel are not major influences on performance, although certainly the two panels offer different coverage of cancer-related genes. We found, as have other groups [28, 12, 15], that different variant callers can disagree widely on the same data. However, we went farther than previous efforts in explaining these differences. First, we found that much can be attributed to various default filtering settings of the algorithms. When harmonized as much as possible, disagreements between algorithms shrink considerably. Importantly, all five algorithms that we tested—MuTect2, SAMtools, VarScan2, Pisces and VarDict—displayed excellent sensitivity for detecting true mutations. The problem is with false positives. Here we found that, using the old bioinformatics strategy of intersecting results, combining results from multiple mutation callers can eliminate many false positives. Other groups have found similar results in variant calling, using simple intersections or more sophisticated combination methods [29, 30]. Importantly, we found that intersecting the results from multiple biological replicates was an even more powerful way of filtering out false positives while maintaining detection sensitivity. Although a costlier option than running multiple algorithms, more research should be done into the tradeoff between accuracy and cost of this strategy, and whether it is relevant to clinical practice.

Our study is not without limitations. We have focused on just five variant callers, chosen on the basis of offering tumour-only mode, and being publicly available, free to use, easy to install, and robustly running. We also limited ourselves to five variant callers in part because Venn diagrams become essentially unreadable when trying to intersect more than five different sets, and we found visualization by Venn diagrams important in understanding the relationships between algorithms. Still, there are other variant callers that could be tested in similar fashion. Another limitation is that the gold-standard variants we wanted the algorithms to identify were either heterozygous or homozygous in comparison with the hg19 human genome. Thus, their allele frequencies were clustered around 50% or 100%. Detecting mutations at lower allele frequencies can be very important, because real tumor specimens are heterogenous with respect to mixing healthy and diseased tissue (particularly in liquid biopsies), and in terms of different cancer subclones [31]. Other studies have used mixtures of DNA from different cell lines to “create” mutations at lower allele frequencies (e.g. [13]), and better probe the relationship between allele frequency and detectability. We currently have similar efforts under way. That being said, some of our heterozygous mutations were coming in at apparent allele frequencies as low as 0.1 or 0.2, particularly when coverage was lower. So we were able to derive some understanding of the relationships between coverage, allele frequency and detectability. Another of our long-term goals is to apply similar analyses to real lung cancer specimens in which gold-standard mutations have been identified, so that we can assess detection rates specifically for cancer-relevant mutations in real patient data. Further, we intend to incorporate those results into broader health economic analyses, going beyond mere accuracy to estimate the value of different sequencing platforms, targeting panels, and bioinformatics pipelines to ultimately improving clinical outcomes.

## Methods

### Cell lines and DNA sequencing data generation

Data was generated from two Coriell cell lines, GM12878 and GM12877. Cells were prepared as standard for targeted clinical sequencing. Briefly, cells were expanded as per the ATCC recommended protocol, harvested, and processed into FFPE cytoblocks. DNA and RNA were isolated for each cell line using minimum of 6 × 10 *µ*m sections and AllPrep DNA/RNA FFPE Kit (Qiagen) according to vendor’s recommended protocol. Nucleic acid quality assessment and quantitation were respectively performed using the Fragment Analyzer (AATI) and Qubit (Thermo Fisher Scientific). One or two sets of biological triplicates were prepared for each cell line. TruSight 170 (Illumina) libraries were prepared using 40 ng DNA or RNA input. Oncomine Focus Libraries (Thermo Fisher Scientific) were prepared using 10 ng DNA or RNA as input. Illumina libraries were sequenced on the NextSeq 500, with 8 RNA and DNA libraries pooled in each High Output 300 cycle run. Oncomine Focus libraries were sequenced on the Ion Torrent, with 6 RNA and DNA libraries pooled on each 318 chip. The Illumina TST170 assay provides full exonic coverage for 170 cancer-associated genes, covering 527,121 total bases, with 3064 genomic intervals. The Oncomine Focus panel covers 29008 total bases in 47 genes, using 269 genomic intervals. In this study, we use only the DNA sequencing data from either panel.

### DNA sequencing data processing

TruSight sequencing data in FASTQ format was quality checked and mapped to the hg19 genome using the recommended BaseSpace pipeline, which relies on Isaac DNA aligner v3.16.02.19. For the Oncomine data, BAM files were obtained from the Ion Torrent online workflow. But because of indexing issues, reads were extracted from the BAM files back into FASTQ format using Picard v2.10.7. Then the reads were mapped to hg19 using BWA aligner version v0.7.17-r1188, producing the BAM files to be used for variant calling. All FASTQ files are available from SRA under accession PRJNA614006.

### Single nucleotide variant calling

We evaluated five publicly available software packages capable of single nucleotide variant (SNV) calling: SAMtools v1.9 [22], VarScan2 v2.3.9 [32], Mutect2 v4.beta.3-SNAPSHOT [24], VarDict [25], and Pisces v5.2.0.1 [26]. All versions were the most recent available at the time of our study. We downloaded all software packages and installed them on our local compute cluster. Each takes as input BAM files of mapped reads or pileups computed from BAM files, e.g. by the mpileup function of SAMtools. Each was used to call variants in tumor-only mode, meaning mutations were called relative to the hg19 genome. Each program outputs some form of variant call file (VCF), from which we extracted SNV calls along with associated information provided by the variant caller, such as depth of coverage, alternative allele frequency, p-value, etc.

Each variant caller has some pre-defined but adjustable parameters, such as: minimum variant frequency, minimum coverage, minimum base quality score and minimum mapping quality score. In our initial tests, we ran each software with its default, recommended parameters. Table 14 summarizes key parameters and their default values for different programs. In later testing, we “harmonized” the parameters of different algorithms to be as similar as possible. Specifically, we set the minimum variant allele frequence to 0.01, the minimum read depth for variant calling to 10 reads, the minimum base call quality to 20, and the minimum read mapping quality to 20.

**Table 14:** Various parameters along with their default values defined in each mutation caller. A dash means that the corresponding parameter was not defined in the caller’s settings.

| Option | MuTect2 | VarScan2 | SAMtools | VarDict | Pisces |
| --- | --- | --- | --- | --- | --- |
| threshold for allele frequency | – | 0.01 | – | 0.05 | 0.01 |
| base quality score threshold | 18 | – | – | – | – |
| max base quality score | – | – | – | – | 100 |
| min map quality | 20 | null | 0 | null | 1 |
| mean map quality | – | null | – | null | – |
| max map quality | null | – | – | – | – |
| min coverage | – | 8 | – | – | 10 |
| max coverage | – | – | 250 | – | – |
| min supporting variant reads | – | 2 | – | – | – |
| min variant quality score | – | – | – | – | 20 |
| strand bias filter | – | – | – | – | 0.5 |
| min reads to strand bias | 2 | – | – | – | – |

### Performance assessment

For the GM12878 and GM12877 cell lines, we obtained gold-standard variant calls from [17]. We assumed that these and only these SNVs should be present in our cell lines. We intersected the SNVs provided by that study with the genomic intervals covered by the TruSight170 or Oncomine Focus panels, to determine which should be detected. If a caller identifies one of these mutations in a particular dataset, it is designated a true positive (TP). If a mutation within the genomic intervals for a panel is not identified by a caller from that dataset, it is designated a false negative (FN). Any called mutation that falls within the genomic regions but that is not on the gold-standard list is considered a false positive (FP). Variant callers are compared based on their TP, FN, FP numbers and scores derived from these, particularly precision = TP / (TP+FP) and sensitivity = TP / (TP+FN).

## Author’s contributions

AK, KT, DJS, PAC, BL, and TJP designed the study. AK, GAP, and TJP analyzed the data. AK, KT, DJS, PAC, BL, and TJP wrote the paper.

## Acknowledgements

The Authors wish to acknowledge the technical support of the Ottawa Hospital Research Institute/University of Ottawa Bioinformatics Core Facility and StemCore Laboratories for their expert technical assistance. This Project is funded by the Government of Canada through Genome Canada and Ontario Genomics, and by the Government of Ontario.

